# Chromosome 7 gain establishes a just-right WNT signaling state during colorectal tumorigenesis

**DOI:** 10.64898/2026.10.12.758507

**Authors:** Irene C. Zaalberg, Paula Cané-Gasull, Martijn Vromans, Meritxell Brunet Guasch, Natália Marčeková, Anna Vinken, Sjors Middelkamp, René Wardenaar, Daniëlle Koot, Beatriz Carvalho, Hugo J. Snippert, Michael D. Nicholson, Floris Foijer, Michael Schubert, Susanne M.A. Lens

## Abstract

Aneuploidy is a hallmark of colorectal cancer, yet the functional consequences of recurrent chromosome gains and losses remain largely unknown. Leveraging advances in human colon organoid culture and chromosome copy number engineering, we here identify the gain of chromosome 7 as an adaptive event following *APC* loss-of-function mutation. Trisomy 7 confers a cell-intrinsic fitness advantage to *APC*-mutant colon organoids, while selective removal of the additional chromosome 7 copy from patient-derived colon cancer organoids severely impairs colorectal cancer organoid growth. By comparing transcriptomes of isogenic disomy 7 and trisomy 7 organoid lines, we found that chromosome 7 gain promotes organoid fitness through attenuation of excessive WNT-driven stemness caused by bi-allelic *APC* mutation. The gain of chromosome 7 thus establishes a ‘just-right’ WNT signaling state. This effect of chromosome 7 gain is conserved from early adenoma-like *APC*-mutant organoids to advanced colorectal cancer organoids carrying additional oncogenic driver mutations. Moreover, consistent with our organoid data, gain of chromosome 7 is particularly common in tumors of colorectal cancer patients whose *APC* genotypes confer relatively high WNT activity. Overall, this work demonstrates that chromosome-specific aneuploidy can act as an adaptive mechanism to optimize, rather than maximize, oncogenic signaling during tumor evolution.

## Introduction

Aneuploidy, defined as an imbalanced number of chromosomes or chromosomal arms, is a defining genomic feature of colorectal cancer (CRC)^1,2^. The majority of colorectal adenomas and carcinomas display numerical chromosomal aberrations, often already detectable at the earliest clinically recognizable stages of tumorigenesis^3–8^. Notably, these karyotypic alterations are non-random: recurrent gains of chromosomes or chromosomal arms 7pq, 8q, 13q and 20q, together with losses of 17p and 18pq, are observed across tumors from independent CRC patients, with individual chromosome arms affected in ∼40–80% of cases^1,2,8–10^. Despite their high prevalence, the mechanisms driving the emergence and persistence of these specific aneuploidies remain poorly understood.

Recurrent aneuploidy patterns such as those observed in colorectal tumors are increasingly thought to reflect context-dependent selective pressures rather than elevated rates of chromosome mis-segregation per se^11,12^. Even with the low chromosome mis-segregation rates typical of healthy tissues (<1%)^13^, the billions of daily cell divisions in the highly proliferative intestinal epithelium likely result in the frequent generation of aneuploid cells^14^. In tumors, mis-segregation rates can rise dramatically, with mis-segregation occurring in up to half of all cell divisions, further increasing karyotypic diversity^15–18^. Yet most aneuploidies decrease cellular fitness, impairing proliferation or triggering cell elimination through apoptosis, immune surveillance, or cell–cell competition^19–21^. Only in permissive (epi)genetic or environmental contexts specific aneuploidies are found to confer a selective advantage, enabling clonal expansion^22^. In fact, experimental evolution studies in yeast and mammalian systems have shown that cells can harness aneuploidy as an adaptive strategy under defined selective pressures, while the same karyotype change remains neutral or harmful in other contexts^23–26^. Whether analogous selection-driven karyotype evolution underlies the recurrent aneuploidies observed in human colorectal tumorigenesis has remained largely untested.

Recent technical advances provide the experimental tools to address this question directly. Human intestinal organoids can be maintained long-term under niche-inspired, semi-physiological conditions that preserve epithelial architecture, differentiation hierarchies, and stem cell function^27–29^. Introduction of defined cancer driver mutations enables the generation of isogenic models representing discrete stages of CRC development and progression^30,31^. Although chromosomal instability and aneuploidy can spontaneously arise in these systems, systematic, chromosome-specific interrogation of aneuploidy has been precluded by the inability to precisely engineer chromosome copy number. Recently developed strategies now overcome this limitation, enabling targeted copy number changes of individual chromosomes in human cells and allowing causal analysis of aneuploidy in controlled, isogenic settings^10,32–36^.

Here, we combine human intestinal organoid technology, *in vitro* evolution, and chromosomal copy number engineering to dissect the origins and functional consequences of aneuploidy during colorectal tumorigenesis. We find that gain of chromosome 7 arises early as an adaptive response to oncogenic WNT hyperactivation and continues to balance WNT activity and stemness throughout CRC progression and tumor maintenance.

## Results

### Recurrent selection of trisomy 7 in *APC*-mutant colon organoids

To investigate the origins of aneuploidy during early colorectal tumorigenesis, we established a genetically defined, isogenic pre-cancerous human organoid model system. The mutation cluster region of *Adenomatous Polyposis Coli (APC)* exon 16 was targeted by CRISPR-Cas9 in healthy tissue-derived human colon organoids from two independent donors (HUB130 and C002N) (Fig 1A, Fig S1A). Bi-allelic disrupting mutations in *APC*, encoding the central negative regulator of WNT signaling, are the near-universal initiating event in microsatellite-stable colorectal tumorigenesis^3^. These mutations confer independence from exogenous WNT-ligand and R-Spondin-1 (RSPO1), two essential niche factors for the maintenance of *APC*-wildtype organoids^28,30^. Indeed, selection under WNT-ligand and RSPO1-depleted conditions yielded multiple surviving organoid clones (Fig S1B), that were expanded into organoid lines.

**Figure 1:**
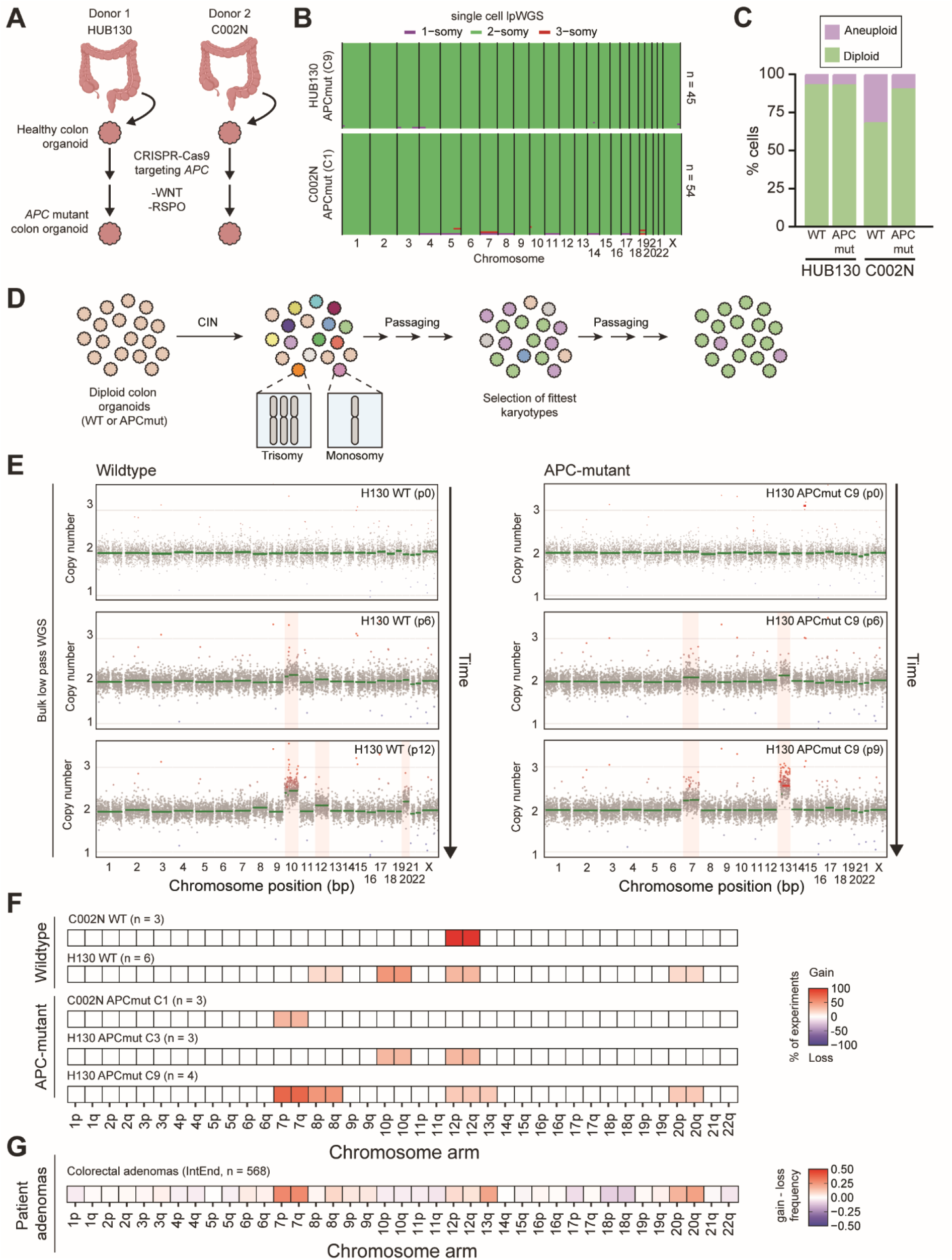
Recurrent selection of trisomy 7 in APC-mutant organoids. A) Strategy for generation of APC-mutant human colon organoids. B) Single cell karyotype-sequencing of HUB130 and C002N APC-mutant organoid lines. C) Quantification of aneuploid cells shown in *Fig 1B*, Fig S1D and Fig S1E. D) Schematic of the in vitro evolution assay: Diploid colon organoids were exposed to MPS1- and CENP-E-inhibitors for 24 hours to generate a population of cells with random aneuploidies. This karyotypically diverse population was subsequently propagated for 12 passages to allow for selection of the fittest karyotypes. E) Examples of bulk lpWGS sequencing profiles of in vitro evolution replicates at indicated passage numbers. Chromosome arms with mean copy number <1.9 or >2.1 are highlighted in red. F) Summary of chromosome arm-copy number alterations detected across all in vitro evolution replicates. G) Gain minus loss frequency (fraction of tumors with gain of indicated chromosome arm minus the fraction of tumors with loss of indicated chromosome arm) of indicated chromosomal arms in human patient colorectal adenomas^8,38^.

Three clonal lines from two different donor backgrounds, with confirmed bi-allelic *APC* disruption (Fig S1C), and with no additional CRC-associated driver alterations (15x WGS, Supplemental file 1), were selected for future experiments. Interestingly, unlike previous reports^37^, the introduced *APC* mutations did not increase intrinsic chromosomal instability (CIN), as evidenced by low aneuploidy and copy number heterogeneity in single cells derived from *APC*-mutant organoid lines (Fig 1B, C, Fig S1D, E). This might stem from a structural distinction in our *APC*-mutant lines: each of our *APC*-mutant clones retained one *APC* allele with an intact C-terminus due to in-frame deletions (Fig S1C), thereby preserving the EB1-binding domain of *APC* which has been implicated in faithful chromosome segregation^37^.

These chromosomally stable *APC*-mutant organoid lines allowed us to interrogate how an *APC*-mutant genetic context influences karyotype evolution. To this end, we performed *in vitro* evolution experiments using the isogenic wildtype and *APC-*mutant organoid pairs. To generate a karyotypically diverse starting population, we first induced a transient pulse of CIN by exposing the organoids to inhibitors of CENP-E (GSK-293295) and MPS1 (NMS-P715) for 24 hours, a regimen that largely preserves chromosome structural integrity and is therefore expected to be better tolerated in the *TP53*-wildtype background of our organoid models^39,40^ (Fig 1D). Single-cell karyotype sequencing confirmed that this approach primarily generated numerical aneuploidies (67-97% of cells), whereas structural aneuploidies occurred at lower frequency (2-18% of cells) (Fig S2). Also, a substantial fraction of cells remained diploid (3-33% of cells) (Fig S2). Following CIN induction, organoid lines were serially propagated for 12 passages. At every third passage, samples were cryopreserved and collected for low pass whole genome sequencing (lpWGS) to monitor karyotype evolution. In total, we performed 9 *in vitro* evolution replicates using wildtype colon organoid lines (3x C002N and 6x HUB130) and 10 replicates using *APC* mutant colon organoid lines (3x C002N APCmut C1, 3x HUB130 APCmut C3, 4x HUB130 APCmut C9), with *APC-*mutant organoids cultured in absence of exogenous WNT-ligand.

Emerging aneuploid organoid populations were detectable in bulk organoid cultures of most *in vitro* evolution replicates, independent of *APC* mutation status (Fig 1E, F, Fig S3). In these bulk cultures, aneuploid copy number states were often non-integer, indicative of sub-clonal aneuploid populations mixed with either diploid cells or cells carrying alternative aneuploidies at low frequency. In replicates wherein aneuploidy was detected, aneuploid copy number states generally increased with prolonged propagation, consistent with positive selection of aneuploid sub-clones (Fig 1E, Fig S3). Endpoint karyotypes were always simple, consisting of one or a few (sub-clonal) whole-chromosome gain(s) (Fig 1E, F, Fig S3). Complex karyotypes and chromosome losses were never detected (Fig 1F), even though they were widespread in the starting population (Fig S2). This indicates counterselection against complex karyotypes and chromosome losses and suggests that only a narrow subset of aneuploid karyotypes is tolerated or beneficial in both healthy and *APC*-mutated genetic contexts.

In wildtype organoids, the most frequently gained chromosomes were 10 and 12, with gains of chromosomes 8 or 20 occurring less frequently (Fig 1F). Similar alterations were also observed in the *APC*-mutant organoid lines (Fig 1F). Strikingly, gain of chromosome 7, the most prevalent early aneuploidy in human colorectal tumorigenesis^8^ (Fig 1G), emerged exclusively and recurrently in the *APC*-mutant genetic context. In HUB130 *APC*-mutant line C9, trisomy 7 emerged in three out of four independent replicates (Fig 1F). Trisomy 7 also emerged sub-clonally in a single replicate of the C002N *APC*-mutant line, although in this case chromosome 7 copy number remained stable at ∼2.1 and did not increase over time (Fig 1F, Fig S3B). Trisomy 7 was never detected in HUB130 *APC*-mutant line C3 (Fig 1F). Trisomy 13, another recurrent early CRC-associated aneuploidy, was detected in a single replicate of HUB130 *APC*-mutant line C9 (Fig 1E, F).

Collectively, these findings demonstrate that relatively simple karyotypic changes, particularly single whole-chromosome gains, can be positively selected in both *APC*-wildtype and *APC*-mutant human colon organoids. Whether the recurrently observed karyotypes shared between wildtype and *APC*-mutant organoids reflect culture-adapted background events or physiologically relevant alterations remains unclear. However, the recurrent emergence of trisomy 7 in a subset of *APC*-mutant organoid lines and patient adenomas suggests that (specific) *APC* mutations may reshape the fitness landscape of early neoplastic cells to favor acquisition of this aneuploidy.

### Trisomy 7 confers a cell-autonomous growth advantage to *APC*-mutant organoids

To interrogate what drives recurrent selection of trisomy 7 in the context of an *APC* mutation, we derived isogenic diploid and trisomy 7 organoid lines from *in vitro*–evolved *APC*-mutant populations of HUB130 C9 described above (Fig 2A). Cryopreserved cultures were reconstituted from passage numbers at which trisomy 7 was present at sub-clonal frequency, as determined by bulk lpWGS. Subcloning single-organoids from these mixed populations yielded multiple clonal, genetically matched *APC*-mutant diploid and trisomy 7 lines (Fig 2B). Unexpectedly, although trisomy 16 was not detected by sequencing of the bulk *in vitro*-evolved population, we identified two *APC*-mutant clones harboring trisomy 16 (Fig 2B). During clonal derivation, a subset of diploid and aneuploid clones acquired *de novo* mutations of unknown significance, as determined by 15x whole genome sequencing (Fig 2C). Importantly, these variants were not universally shared among clones, and we identified clones without *de novo* mutations for each karyotype (Fig 2C). This established an experimental system in which the phenotypic effects of trisomy 7 could be interrogated independently of confounding secondary genetic events.

**Figure 2:**
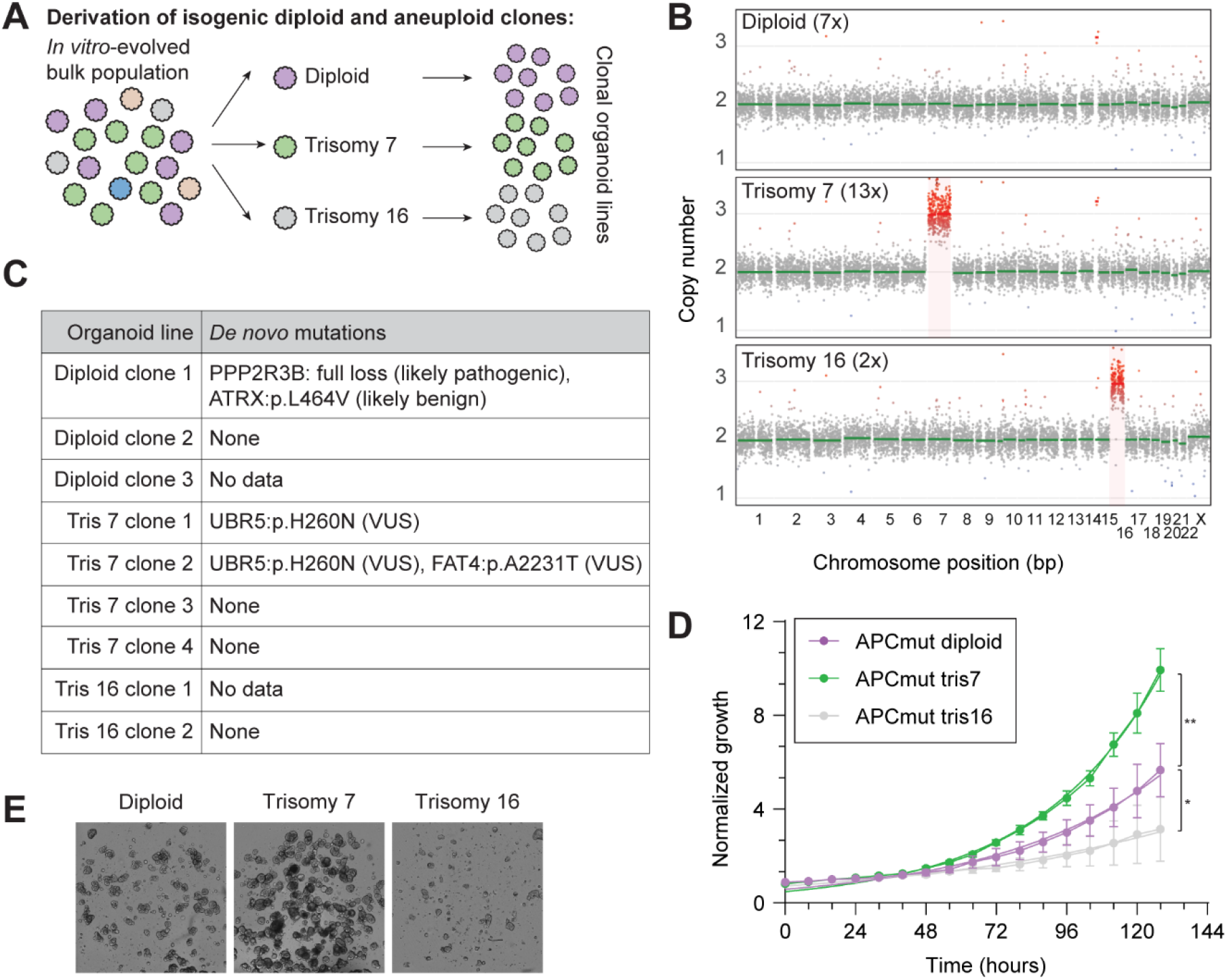
Trisomy 7 confers a cell-autonomous growth advantage. A) Strategy for derivation of isogenic diploid and aneuploid organoid lines by single-organoid cloning: cryopreserved organoids harboring sub-clonal chromosome 7 gain (copy number between 2 and 3) were thawed and single-cell plated. Individual organoids were isolated and expanded clonally. B) Representative bulk lpWGS profiles of representative clonal organoid lines derived as described in A. C) Clone-specific mutations identified by 15x whole genome sequencing. D) Representative IncuCyte growth curves of APC-mutant diploid, trisomy 7, and trisomy 16 organoids. Data are presented as mean ± SD and are based on two independent organoid lines per genotype. Growth rates were compared using nonlinear regression and extra-sum-of-squares F tests (* = p < 0.05. ** = p < 0.005). E) Representative endpoint images corresponding to D.

Positive selection of trisomy 7 could theoretically arise from either a cell-intrinsic proliferative advantage or competitive interactions with neighboring diploid cells. To directly test for cell-autonomous fitness effects, we quantified proliferation using multiple orthogonal assays. Across all assays, trisomy 7 organoids exhibited a significant increase in proliferative capacity relative to their diploid counterparts (Fig 2D, E, Fig S4A, B). By contrast, trisomy 16 organoids displayed impaired proliferation. In fact, the reduced proliferative capacity of the trisomy 16 clones affected material availability to such an extent, that we were unable to include both clones in all experimental replicates. These results demonstrate that trisomy 7 confers a chromosome-specific, cell-autonomous proliferative advantage in an *APC*-mutant genetic background. This intrinsic fitness gain provides a mechanistic explanation for its recurrent positive selection during early colorectal tumorigenesis, while not excluding potential additional contributions from competitive interactions.

### Trisomy 7 addiction in established colorectal cancer

We next asked whether trisomy 7 remained relevant for tumor cell fitness at later stages of CRC progression, following the accumulation of additional oncogenic driver mutations. To this end, we aimed to reduce chromosome 7 copy number from three to two in the P9T patient-derived organoid line, an established microsatellite-stable (MSS) and chromosomally instable CRC model^16,41,42^. P9T harbors mutations in the four most common CRC driver genes (*APC*, *TP53*, *KRAS*, and *PIK3CA*) alongside various aneuploidies, including three copies of chromosome 7^16,41,42^ (Fig 3A).

**Figure 3:**
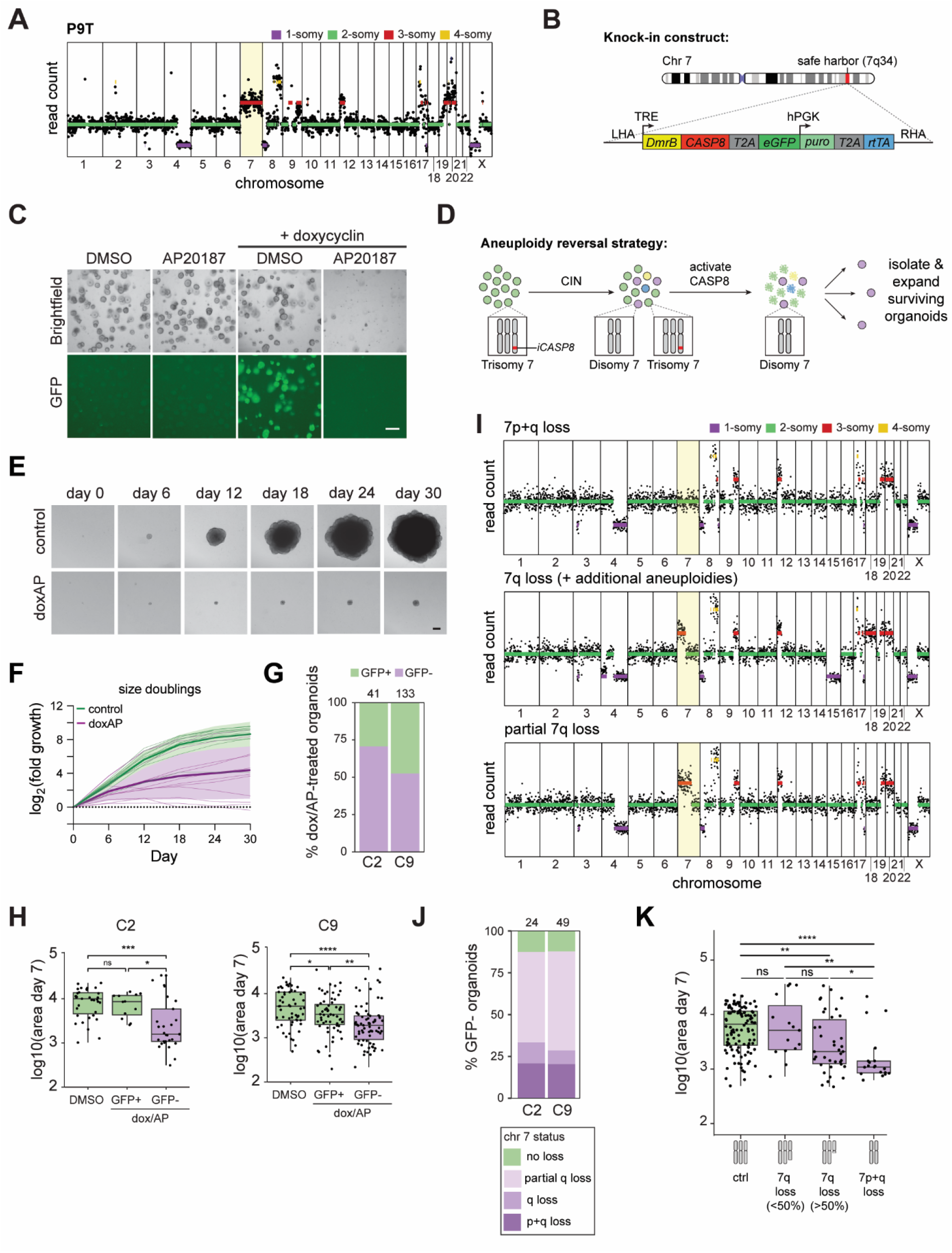
Trisomy 7 to disomy 7 reversal impairs growth of colorectal carcinoma organoids. A) Bulk lpWGS profile of P9T CRC organoids. B) Schematic of the iCASP8-eGFP-puro-based selection cassette integrated into a safe-harbor locus at chromosome 7q34. C) Brightfield and GFP images of P9T CRC chromosome 7-iCASP8 organoids treated with DMSO, AP20187 (AP), doxycyclin (dox)/DMSO or dox/AP. Scale bar, 200 µm. D) Strategy to enrich for cells that lost the iCASP8-tagged chromosome 7 copy through spontaneous chromosome mis-segregation. E) Representative images of control and doxycycline/AP20187-surviving organoids after prolonged culture. Scale bar, 200 µm. F) Quantification of organoid growth over 30 days following single-organoid isolation (n = 13 control and n = 12 dox/AP-recovered organoids). Thick lines indicate the mean, thin lines represent individual organoids, shaded areas represent standard deviation. G) Percentage of dox/AP-recovered organoids classified as GFP-positive or GFP-negative after doxycycline re-exposure. H) Projected organoid area at day 7 after single-organoid isolation for two independent clonal chromosome 7-iCASP8 P9T CRC organoid lines. Box plots show the median (horizontal line), interquartile range (box) and 1.5 × IQR. Dots represent individual organoids. Two-sided Wilcoxon rank-sum test, * = p < 0.05, ** = p < 0.005, *** = p < 0.0005, **** = p < 0.00005. GFP status was determined by doxycyclin re-exposure at > 7 days post-isolation. I) Representative lpWGS profiles of GFP-negative organoids recovered after dox/AP selection. J) Distribution of chromosome 7 copy-number states among GFP-negative, dox/AP-recovered organoids. K) Projected organoid area at day 7 after single-organoid isolation for dox/AP-recovered organoids, stratified by karyotype. Box plots show the median (horizontal line), interquartile range (box) and 1.5 × IQR. Dots represent individual organoids. Two-sided Wilcoxon rank-sum test, * = p < 0.05, ** = p < 0.005, **** = p < 0.00005.

To selectively eliminate one copy of chromosome 7, we introduced an inducible CASPASE8 (iCASP8)-based negative selection cassette into the 7q34 safe-harbor locus using CRISPR-Cas9 technology^43–45^ (Fig 3B). Multiple independent knock-in clones were validated for correct mono-allelic integration at 7q34 by genomic PCR and quantitative PCR (Fig S5, Methods). In this system, doxycyclin (dox) induces expression of DmrB (FKBP-F36V)-CASP8 together with eGFP, while additional supplementation of AP20187 (AP) promotes DmrB-mediated CASP8 dimerization and apoptosis (Fig 3C). Combined dox/AP treatment should selectively eliminate cells carrying the cassette-bearing chromosome 7 copy and thus enrich for cells that lost (part of) this chromosome through previous CIN events (Fig 3D).

To derive isogenic trisomy-to-disomy-7-reversed organoid lines, we isolated and expanded the few organoids that survived dox/AP selection (Fig 3D). Using projected organoid area as a proxy for organoid size, we noted that clonal outgrowth of many GFP-negative dox/AP-recovered organoids was severely impaired compared to identically processed controls (Fig 3E-H). Of note, a substantial fraction of dox/AP-recovered organoids remained GFP-positive, suggesting that they acquired dox/AP resistance through other mechanisms than trisomy 7 loss (Fig 3G). In contrast to the GFP-negative population, these GFP-positive organoids showed no or a modest growth impairment (Fig 3H). LpWGS confirmed that chromosome 7 copy number alterations were present in the majority of GFP-negative organoids (87.7% of 73 analyzed GFP-negative dox/AP-recovered organoids), whereas GFP-positive organoids had in most cases retained three chromosome 7 copies (87.5% of 12 analyzed GFP-positive dox/AP recovered organoids) (Fig 3J, Fig S6C). Therefore, GFP-positive organoids were excluded from further analyses.

Most copy number alterations in the GFP-negative dox/AP recovered organoids (57.5% of all organoids) consisted of partial, terminal 7q deletions encompassing the iCASP8-casette at 7q34 (Fig 3I, bottom, Fig 3J). Loss of the entire 7q arm was detected in 9.6%, and loss of a full copy of chromosome 7pq in 20.5% of dox/AP-surviving organoid clones (Fig 3J). Strikingly, organoids that lost a full copy of chromosome 7 exhibited the most severe proliferation defects (Fig 3K). Organoids that lost >50% of their third 7q arm also exhibited reduced growth, albeit less pronounced than that observed in organoids with complete chromosome 7 loss (Fig 3K). In contrast, organoids that lost <50% of their third 7q arm did not show a significant reduction in growth, providing an internal control for potential growth effects of the reverse engineering procedure itself (Fig 3K). Thus, reversal to chromosome 7 disomy causes a growth defect in established CRC organoids, with the severity of the growth impairment depending on the extent of chromosome 7 copy number loss.

Together, these findings demonstrate that trisomy 7 is not merely an early, transient adaptation but conveys a sustained contribution to tumor cell fitness in established CRC despite the accumulation of additional oncogenic driver alterations. The marked proliferative impairment following loss of the supernumerary chromosome 7 suggests that progressed CRC cells become ‘addicted’ to this aneuploidy^33^, and highlights trisomy 7 as a potential vulnerability across disease stages

### Trisomy 7 is associated with reduced WNT signaling and stemness

To elucidate how chromosome 7 gain promotes tumor growth, we compared transcriptomes of isogenic trisomy 7 and disomy 7 organoid models. In the *APC*-mutant setting, we performed bulk RNA sequencing on two independent trisomy 7 clones, a matched diploid clone derived from the same bulk culture, and the parental diploid population (Fig 4A). For the CRC model, we implemented a single-organoid RNA sequencing approach, to allow for selective inclusion of organoids with 7q or 7pq loss. For this, size-matched organoids, derived from untreated or dox/AP-treated P9T chromosome 7-iCASP8 cultures, were collected (Fig 4A, Fig S7A, B). Chromosomal copy number was inferred from transcriptomic data using an expression-based karyotyping strategy (see Methods). The robustness of this approach was supported by down-sampling analyses of bulk RNA-seq datasets and orthogonal validation using matched lpWGS (Fig S7C, D).

**Figure 4:**
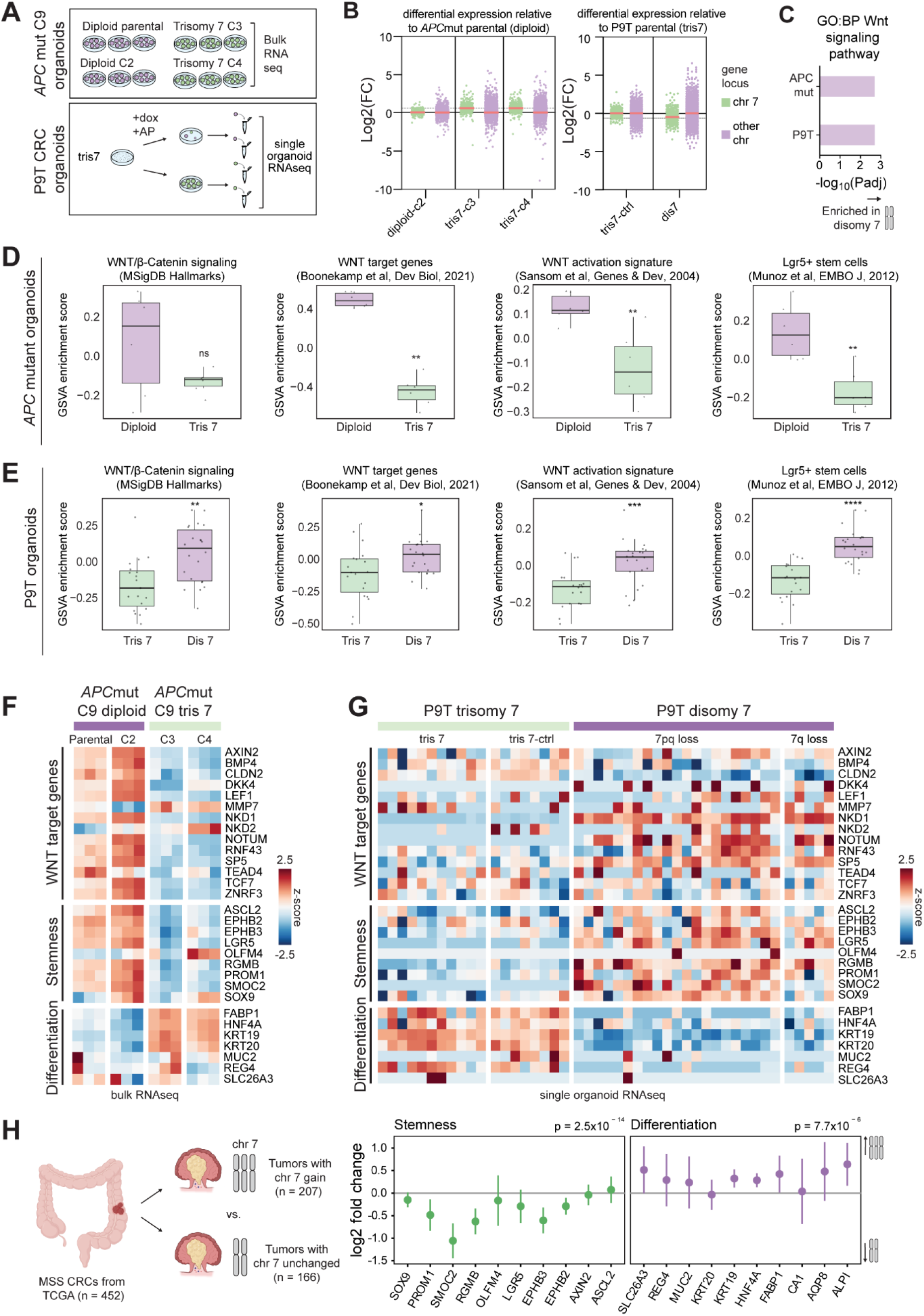
Trisomy 7 is associated with reduced WNT signaling and stemness. A) RNA-sequencing strategy. B) Differential expression of chromosome 7-encoded genes relative to genes on all other chromosomes in isogenic disomy 7 and trisomy 7 organoid lines, normalized to their parental organoid line. C) Gene ontology enrichment analysis of trisomy 7 versus disomy 7 organoids, in P9T CRC and APC-mutant organoid lines. D) Gene set variation analysis (GSVA) of published WNT signaling and stemness signatures in APC-mutant diploid and trisomy 7 organoid lines^48–50^. Box plots show the median (horizontal line), interquartile range (box) and 1.5 × IQR. Dots represent individual organoids. Dots represent 3 technical replicates of 2 independent organoid lines. Two-sided Wilcoxon rank-sum test, ** = p < 0.005. E) GSVA of published WNT signaling and stemness signatures in P9T trisomy 7 and disomy 7 organoids^48–50^. Box plots show the median (horizontal line), interquartile range (box) and 1.5 × IQR. Dots represent individual organoids. Dots represent individual organoids. The disomy 7 category includes a minority of organoids with 7q loss only, to increase sample size and statistical power. Two-sided wilcoxon rank-sum test, * = p < 0.05, ** = p < 0.005, *** = p < 0.0005, **** = p < 0.00005. F, G) Expression heatmap of core WNT signaling, stemness and differentiation genes. H) Expression of stemness and differentiation genes in microsatellite stable, POLε-wildtype colorectal tumors from TCGA, with or without gain of chromosome 7, adjusted for APC alteration, whole genome duplication, TP53 mutation, RAS/RAF mutation, and tumor site (Wald test; 95% CIs calculated using Wald standard errors; gene-level Wald p-values combined using Fisher’s method).

Consistent with previously described gene dosage effects^46,47^, trisomy 7 was associated with increased expression of chromosome 7-encoded genes (Fig 4B): gain of trisomy 7 in *APC*-mutant organoids resulted in an average 48% increase in gene expression, and loss of trisomy 7 in CRC organoids in an average 30% decrease. Thus, RNA expression changed near-linearly with DNA copy number (an expected 50% increase or 33% decrease, respectively).

Gene ontology analysis identified a common transcriptional change associated with trisomy 7 across organoid models, namely decreased WNT signaling in *APC*-mutant organoids following chromosome 7 gain, and increased WNT signaling in CRC organoids following trisomy 7 to disomy 7 reversal (Fig 4C). Further interrogation of our RNA-sequencing data confirmed this finding: trisomy 7 was associated with a reduction of WNT-signaling associated genes in *APC*-mutant organoids (Fig 4D, F)^48–50^. This decrease was accompanied by reduced expression of canonical stem cell markers, including *LGR5*, *ASCL2*, *PROM1*, and *SMOC2*, alongside increased expression of differentiation-associated genes such as *KRT20* and *FABP1* (Fig 4F)^48–50^. This inverse relationship between chromosome 7 dosage and WNT-driven stemness was recapitulated in the CRC model: organoids that lost their third copy of chromosome 7pq or 7q displayed marked enrichment of WNT signaling and stemness programs (Fig 4E, G). To determine whether this relationship was also observed in human tumors, we analyzed RNA-sequencing data from *APC*-mutant CRCs in The Cancer Genome Atlas (TCGA) database^2^. Consistent with our organoid data, tumors harboring a chromosome 7 gain displayed reduced expression of stemness-associated genes and relative enrichment of differentiation markers compared to tumors in which chromosome 7 copy number was unchanged (Fig 4H).

Collectively, these findings support a functional interplay between chromosome 7 gain and WNT signaling and stemness regulation in CRC.

### Trisomy 7 restrains WNT-driven stemness

At first glance, the association between trisomy 7 and reduced WNT signaling appears paradoxical. Constitutive WNT pathway activation due to APC loss is a key initiating event in colorectal tumorigenesis, enabling sustained proliferation independent of the WNT niche. However, accumulating evidence suggests that tumor growth requires finely tuned, ‘just-right’ levels of WNT pathway activity, with excessive signaling imposing a proliferative constraint^51,52^. Consistent with this concept, healthy colon organoids displayed a clear proliferative optimum in response to the WNT pathway activator CHIR at concentrations close to 2 µM when cultured in absence of exogenous WNT-ligand (Fig 5A, B). Both lower and higher CHIR concentrations markedly impaired growth. In contrast, CHIR concentrations as low as 0.5 µM already impaired growth of *APC*-mutant HUB130 and P9T organoids, which proliferated maximally in the absence of CHIR (Fig 5A–C), indicating that basal WNT signaling in these lines meets or exceeds the proliferative optimum. In line with this interpretation, partial WNT suppression using a low dose of the β-catenin-TCF inhibitor iCRT3, sufficient to reduce *LGR5* expression (Fig 5D), increased the proliferation of *APC*-mutant diploid organoids (Fig 5E). This effect was not observed in *APC*-mutant or CRC organoids harboring trisomy 7 (Fig 5E, F). Moreover, the dose at which iCRT3 treatment switched from neutral or growth-promoting to growth-inhibitory occurred at lower concentrations in *APC*-mutant organoids with trisomy 7 than diploid *APC* mutant organoids, consistent with trisomy 7 lowering basal WNT pathway activity (Fig 5E).

**Figure 5:**
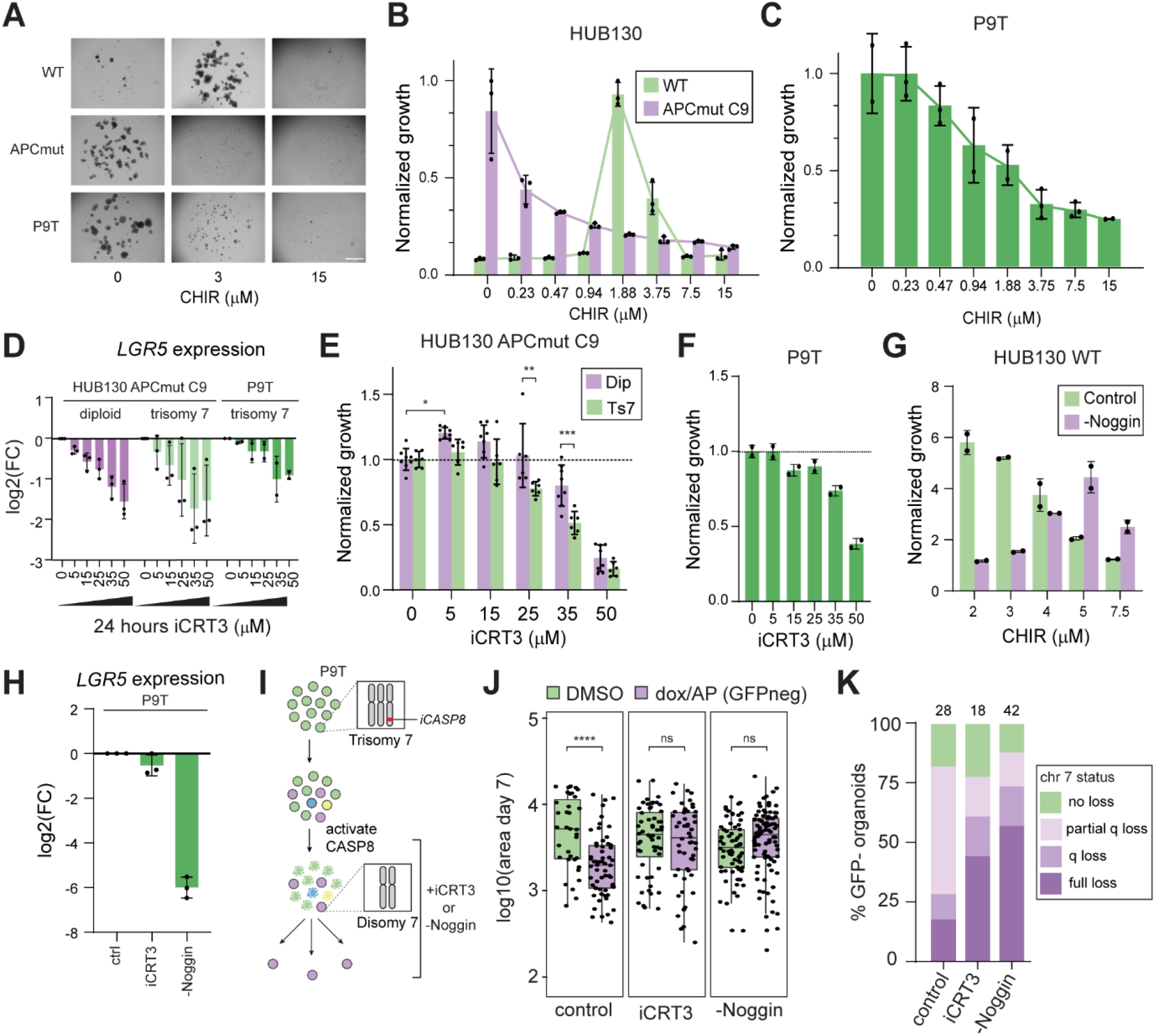
Trisomy 7 restrains WNT-driven stemness to optimize proliferative fitness. A) Brightfield images of APC-wildtype organoids, APC-mutant organoids and P9T CRC organoids cultured in increasing concentrations of the WNT activator CHIR99021 (in absence of exogenous WNT-ligand). B, C) IncuCyte-based growth responses to CHIR99021 in HUB130 wildtype, HUB130 APC-mutant C9 (B) and P9T (C) organoids. Mean ±SD. Dots represent technical replicates. D) LGR5 expression in HUB130 wildtype, HUB130 APC-mutant C9 and P9T organoids treated with increasing concentrations of β-catenin-TCF inhibitor iCRT3 (RTqPCR). Mean ±SD. Dots represent biological replicates. E) IncuCyte-based growth responses to increasing concentrations of β-catenin-TCF inhibitor iCRT3 in HUB130 APC-mutant diploid and APC-mutant trisomy 7 organoids. Mean ±SD. Dots represent biological replicates and are derived from 2 independent organoid lines per genotype. Two-way ANOVA, * = p < 0.05, ** = p < 0.005, *** = p < 0.0005. Only selected relevant pairwise comparisons are shown for clarity. F) Representative IncuCyte-based growth responses to increasing concentrations of β-catenin-TCF inhibitor iCRT3 in P9T organoids. Mean ±SD. Dots represent technical replicates. G) Representative IncuCyte-based growth response of HUB130 wildtype organoids cultured with increasing CHIR99021 concentrations in presence or absence of Noggin (in absence of WNT-surrogate). Mean ±SD. Dots represent technical replicates. H) LGR5 expression following treatment with 15 µM iCRT3 or Noggin withdrawal in P9T organoids (RTqPCR). Mean ±SD. Dots represent biological replicates. I) Schematic of disomy 7-rescue experiments. J) Projected organoid area of dox/AP-treatment surviving P9T organoids at 7 days post single organoid isolation following treatment with 15 µM iCRT3 or Noggin withdrawal. Box plots show the median (horizontal line), interquartile range (box) and 1.5 × IQR (whiskers). Dots represent individual organoids. K) Chromosome 7 copy number status in dox/AP-surviving organoids shown in J.

These observations prompted us to test whether attenuating WNT signaling or reducing stemness could mitigate the growth defect associated with trisomy 7 to disomy 7 reversal in CRC organoids. To dampen WNT pathway activity while avoiding toxicity, P9T organoids were treated with a tolerated dose of 15 µM iCRT3. As an alternative way to reduce stemness, we withdrew Noggin from the culture medium. Noggin is a BMP-signaling antagonist that promotes stemness by hampering BMP-mediated differentiation^53^. In our *APC*-wildtype colon organoids, removal of Noggin from the culture medium increased the growth-optimal dose of CHIR from 2 µM to 5 µM (Fig 5G), indicating that BMP signaling negatively regulates WNT signaling. In P9T CRC organoids, both iCRT3 treatment and Noggin removal reduced stemness, as assessed by *LGR5* expression, with Noggin withdrawal exerting a substantially stronger effect than iCRT3 treatment (Fig 5D, H). Strikingly, both treatments rescued the growth defect of GFP-negative organoids recovered following dox/AP-treatment of our Chr7-iCASP8 P9T organoid line (Fig 5I, J). Moreover, the percentage of recovered organoids with a full chromosome 7 loss was markedly increased (Fig 5K). Possibly, under standard culture conditions many disomy 7 cells either die or fail to develop into organoids, resulting in their underrepresentation in the recovered population following dox/AP treatment. Attenuation of WNT signaling and stemness programs appears to mitigate these constraints, enabling survival and expansion of cells with a complete trisomy-to-disomy 7 reversion. More broadly, these findings identify elevated WNT-driven stemness as causal determinant of the fitness barrier imposed by trisomy 7 loss in CRC organoids.

### Differences in *APC* genotype affect the selective pressure for chromosome 7 gain

In CRC, bi-allelic *APC* inactivation typically occurs through truncating mutations. Such mutations eliminate the C-terminal region of the protein containing a series of 20-amino acid repeats (20Rs), which are required for efficient β-catenin binding and degradation. The number of 20Rs retained across both mutant alleles therefore provides a quantitative readout of the residual *APC*-mediated restraint on WNT signaling: mutations that eliminate all 20Rs result in maximal WNT pathway activation, whereas mutations retaining one or more 20Rs preserve partial β-catenin degradation capacity and thus lead to submaximal WNT activation^54,55^. As the organoid data demonstrated that the fitness benefit of chromosome 7 copy number gain specifically manifests in the context of elevated WNT-driven stemness, we hypothesized that chromosome 7 gain would be preferentially selected in the context of *APC* genotypes conferring relatively high WNT activity.

Such a model could explain the divergent evolutionary trajectories of the *APC*-mutant HUB130 organoid lines C3 and C9, in which trisomy 7 was repeatedly acquired by C9 but never by C3 during *in vitro* evolution (Fig 1F). Indeed, although both clones harbored a similar truncating mutation in one *APC* allele, the second allele differed. In C9, the second *APC* allele lacked the region encoding the second and third 20-amino acid repeats (20R2 and 20R3), including the β-catenin inhibitory domain (CID), whereas C3 retained both the CID and the 20R3 repeat (Fig 6A). Although the large CRISPR-induced deletions in these clones do not precisely recapitulate the typical truncating *APC* mutations observed in human tumors, the CID and 20R3 are considered particularly important determinants of APC-mediated β-catenin degradation, predicting greater constitutive WNT activity in C9 than in C3^54,56,57^. Functional analyses supported this prediction: C3 was less responsive to CHIR-mediated WNT hyperactivation than C9, indicative of lower basal WNT pathway activity in C3 (Fig 6B). Furthermore, C9, but not C3, proliferated in the absence of Noggin (Fig 6C). Whole-genome sequencing identified no oncogenic genetic alterations affecting BMP pathway components (Supplemental file 1), indicating that the Noggin-independent phenotype of C9 is unlikely to result from BMP pathway mutations. Instead, it suggests that C9 possesses a greater intrinsic capacity to maintain stemness, most likely as a consequence of more elevated basal WNT signaling due to its more severe APC mutational genotype.

**Figure 6:**
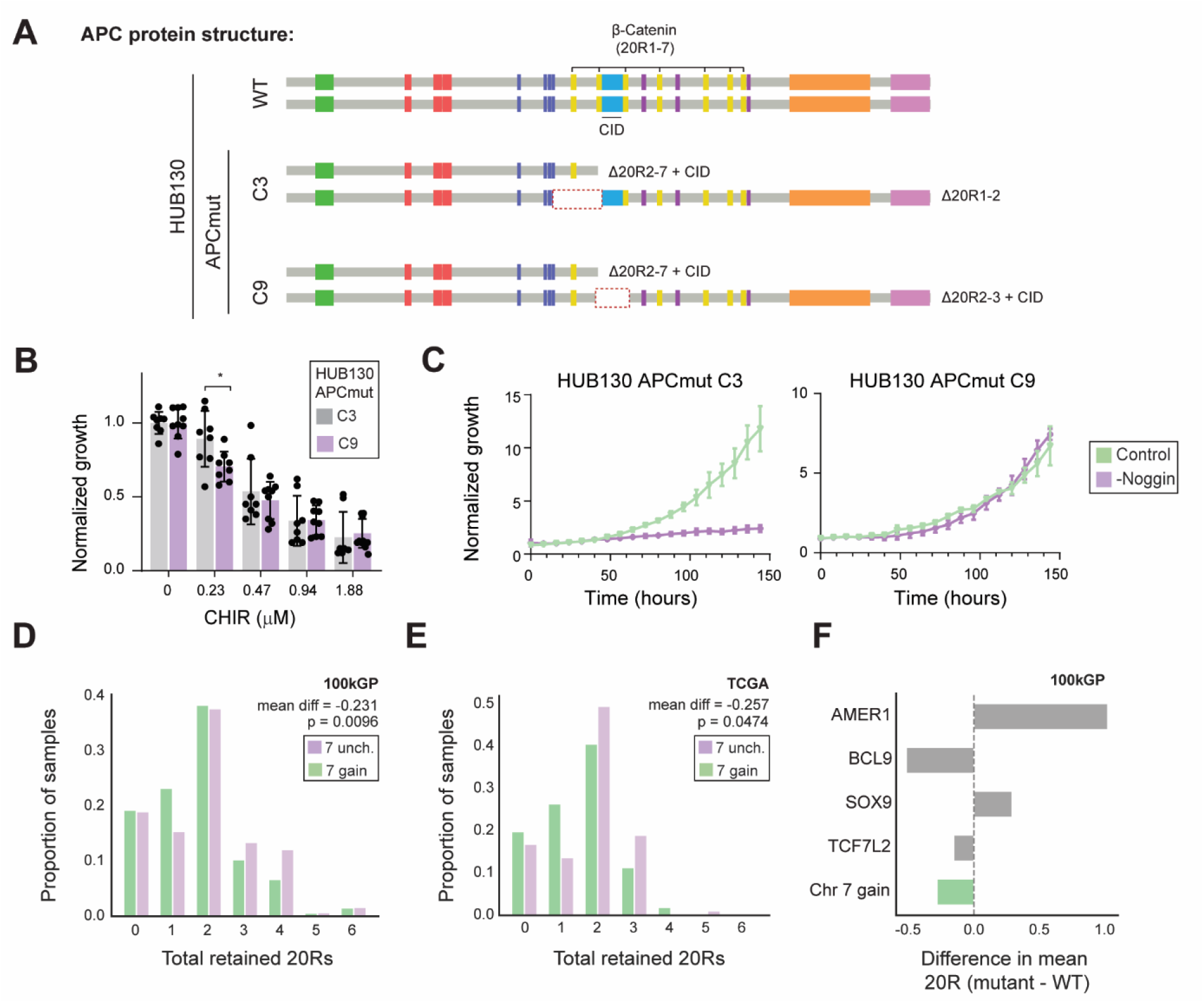
Differences in APC genotype affect selective pressure for trisomy 7. A) Predicted APC protein structures in HUB130 wildtype, APC-mutant C3 and APC-mutant C9, inferred from DNA sequence (Fig S1C). B) IncuCyte-based growth of HUB130 APC-mutant C3 and HUB130 APC-mutant C9 organoids cultured in increasing CHIR99021 concentrations. Mean ± SD. Dots represent technical replicates derived from three independent experiments. Two-sided Wilcoxon rank-sum test, * = p < 0.05. C) IncuCyte-based growth curves of HUB130 APC-mutant C3 and HUB130 APC-mutant C9 organoids cultured in absence or presence of Noggin. D) Distribution of total retained APC 20Rs in primary, microsatellite-stable colorectal tumors from the 100kGP database stratified by chromosome 7 status. Chromosome 7 gain, n = 524. Chromosome 7 unchanged, n = 305. Mean difference = −0.231. Mann-Whitney p-value = 0.00962. E) Distribution of total retained APC 20Rs in primary, microsatellite-stable colorectal tumors from the TCGA database stratified by chromosome 7 status. Chromosome 7 gain, n = 106. Chromosome 7 unchanged, n = 95. Mean difference = −0.257. Mann-Whitney p-value = 0.0474. F) Difference in mean total retained 20Rs (mutant - unchanged) in the 100kGP for tumors carrying oncogenic mutations in the WNT regulators AMER1, BCL9, SOX9, or TCF7L2, shown alongside chromosome 7 gain for comparison. Bars to the left of zero indicate that the alteration is associated with APC genotypes with more complete APC inactivation / higher WNT activity relative to tumors without the mutation, and vice versa.

We then asked whether variation in *APC* genotype was associated with chromosome 7 gain in patient tumors. If higher basal WNT activity increases the selective advantage of chromosome 7 gain, we reasoned that tumors harboring chromosome 7 gain should carry on average a lower total number of retained 20Rs than tumors without chromosome 7 copy-number alteration. To test this prediction, we surveyed human colorectal primary tumors in both the 100,000 Genomes Project (100kGP) and TCGA cohorts. In both cohorts, microsatellite-stable primary CRCs with a chromosome 7 gain carried significantly fewer total retained 20Rs than tumors without the alteration (Fig 6D, E). The difference in mean 20R of tumors with chromosome 7 gain fell within the range previously observed for secondary regulators of the “just-right” WNT optimum^55^, such as *BCL9* or *TCF7L2* mutations (Fig 6F).

Together, these observations suggest that the selective advantage conferred by trisomy 7 depends on the pre-existing WNT signaling landscape in colorectal tumors, with chromosome 7 gain selectively favored in tumors with relatively high WNT activity.

## Discussion

Colorectal tumorigenesis is characterized by recurrent genetic alterations, including frequent chromosome-level copy number changes. Among these, chromosome 7 gain is one of the earliest and most recurrent events^8^, yet whether it represents a tolerated consequence of chromosomal instability or an actively selected driver has remained unresolved. Here, using genetically matched organoid models, we show that chromosome 7 gain is not simply tolerated but confers a context-specific selective advantage by attenuating WNT pathway hyperactivation following *APC* mutation. These findings explain the recurrent gain of chromosome 7 in CRC and establish this early chromosome-specific aneuploidy as an active contributor to colorectal tumorigenesis.

Our findings support and extend the emerging concept that cancer cells evolve to maintain oncogenic signaling within a permissive ‘Goldilocks’ zone, in which pathway activity is sufficient to sustain proliferation but constrained below the threshold at which excessive signaling compromises cellular fitness^58–60^. While this principle has been recognized for WNT signaling through the ‘just-right’ model of allelic *APC* mutation selection and secondary alterations in WNT pathway components such as *TCF7L2* and *BCL9*^51,55,58,61^, we identify chromosome 7 gain as an additional mechanism by which colorectal tumor cells tune WNT signaling toward this optimal fitness window. A conceptually related example comes from Ewing sarcoma, where recurrent gain of chromosome 8, through increased dosage of cohesin complex-component *RAD21*, dampens replication stress associated with oncogenic EWS-FLI1 fusion^62^. Together, these findings suggest that specific aneuploidies may be selected not simply to amplify oncogenic driver gene output, but to establish cellular states that optimize the trade-off between oncogenic activity and its associated fitness costs.

A central implication of the optimal WNT signaling model is that the fitness effect of chromosome 7 gain depends on the pre-existing signaling state of the cell. Consistent with this principle, trisomy 7 was not selected in every *APC*-mutant clone or across every independent replicate. This variability mirrors human colorectal adenomas, in which chromosome 7 gain is among the earliest and most frequent chromosomal alterations, but is not universally observed^8^. We propose that this variability in chromosome 7 gain likelihood reflects differences in the baseline WNT signaling state established by the initiating *APC* mutations. Different *APC* mutations generate distinct levels of residual β-catenin and consequently different WNT signaling outputs^54,55,63^, creating different selective pressures for subsequent pathway modulation. Within this framework, chromosome 7 gain would be selected only when it shifts WNT signaling closer to the optimal fitness window (Fig 7). Consistent with this model, patient tumors with fewer retained *APC* 20Rs, associated with higher WNT pathway activation, were more likely to harbor chromosome 7 gain. Chromosome 7 gain should therefore not be viewed as an inevitable consequence of *APC* loss, but as one of several evolutionary solutions that enable colorectal tumors to achieve an optimal level of WNT pathway activity and thereby maximize cellular fitness.

**Figure 7:**
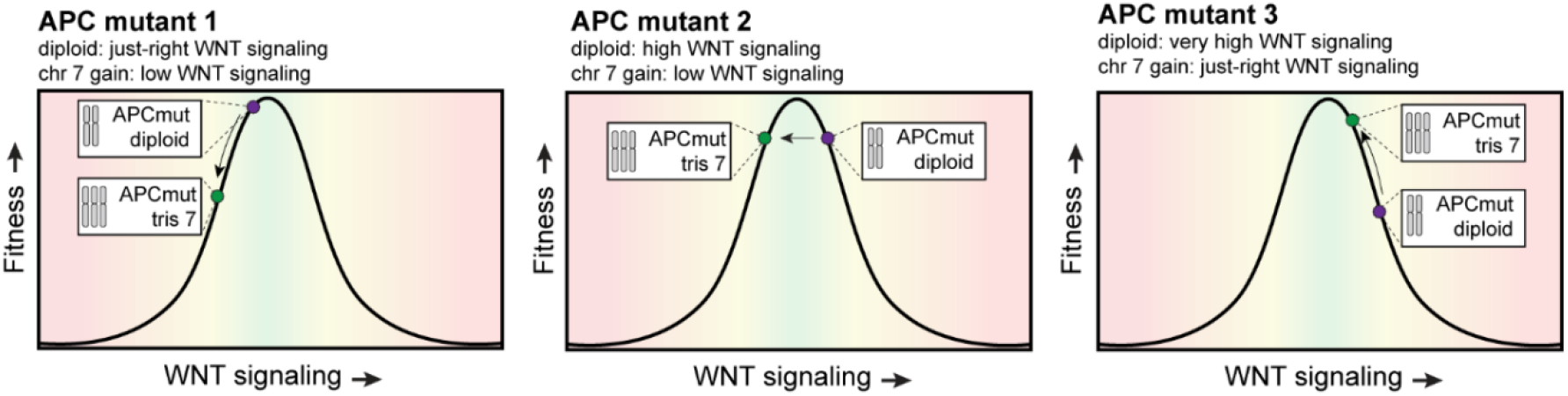
Model for the context-dependent fitness effects of chromosome 7 gain. APC mutations determine the baseline level of WNT signaling, with maximal fitness is at an intermediate level of pathway activity. Chromosome 7 gain reduces WNT signaling and therefore has distinct effects depending on the initial signaling state. When WNT activity is low or near the fitness optimum, chromosome 7 gain is detrimental or neural (APC mutants 1 and 2, respectively). When WNT activity is excessive, however, chromosome 7 gain shifts WNT signaling toward the optimum and confers a fitness advantage (APC mutant 3). Colored dots indicate the predicted WNT signaling and fitness states of diploid and trisomy 7 cells.

How chromosome 7 gain restrains WNT signaling and stemness remains unclear. Given that the impact of chromosome 7 loss was proportional to the extent of chromosomal material lost, it is likely that multiple dosage-sensitive genes contribute cumulatively to the fitness advantage conferred by trisomy 7, rather than a single dominant driver. Interesting candidates to contribute to reduced WNT signaling following chromosome 7 gain include *HOXA5* (7p), which suppresses intestinal stem cell programs and antagonizes WNT signaling in colorectal cancer^64^, and *CAV1* (7q), which has been implicated in restricting β-catenin transcriptional activity by regulating its subcellular localization^65–67^. *CDK6* (7q) has been proposed to promote β-catenin degradation through phosphorylation, potentially limiting WNT transcriptional output^68^. Other chromosome 7 genes, including *BCL7B* (7q) and *DBF4* (7q), are potential modulators of β-catenin-dependent transcription through effects on chromatin accessibility at β-catenin-responsive target genes^69–71^. An alternative, and potentially complementary, mechanism is that increased dosage of chromosome 7-encoded MAPK activators, including *BRAF*, *EGFR* and *MET*, enhances MAPK signaling, thereby attenuating WNT pathway output and preventing excessive stem cell activity. MAPK and WNT signaling pathways have been described to engage in extensive crosstalk in the colorectal epithelium, with MAPK activity constraining β-catenin–dependent transcriptional programs^72–76^. Moreover, inhibition of MAPK regulators, including *EGFR*, initiates a WNT-high, stem-like cell state in colorectal (cancer) epithelium, characterized by high expression of LGR5^72–74,76,77^. Such a LGR5-high cell state is associated with quiescence and reduced proliferative output^72,78^, reminiscent of the phenotype that we observe following trisomy 7 to disomy 7 reversal in P9T CRC organoids. Future experiments that selectively perturb dosage or activity of these candidate MAPK and WNT regulators will be important to determine whether MAPK–WNT crosstalk contributes to the effects of trisomy 7, and to establish whether this represents a major mechanism through which chromosome 7 gain promotes cellular fitness.

Remarkably, trisomy 7 remained indispensable for advanced CRC organoids carrying additional driver mutations in *KRAS*, *PIK3CA*, and *TP53*. Rather than acting as a temporal surrogate for individual oncogenic mutations, chromosome 7 gain may establish a signaling and transcriptional state that cannot be fully recreated by subsequent genetic alterations. Finally, the extra copy of chromosome 7 may become progressively integrated into the evolving cancer genome^79^. In this way, analogous to the concept of oncogene addiction, and as previously proposed, tumors may develop a dependence on specific aneuploidies as they evolve^33^. As additional genetic and epigenetic alterations accumulate, cancer cells progressively adapt to the altered cellular state imposed by trisomy 7, resulting in an increasing reliance on the aneuploid state itself. Such evolutionary path dependence provides a potential explanation for why trisomy 7 not only persists during CRC progression but also becomes increasingly difficult to eliminate without compromising tumor fitness.

Overall, our study provides a mechanistic explanation for one of the earliest and most recurrent chromosome-level alterations in colorectal cancer. Trisomy 7 is actively selected following *APC* mutation because it optimizes WNT signaling output, thereby promoting proliferation and establishing a signaling state that remains advantageous throughout tumor progression. Hence, we identify aneuploidy as a mechanism to tune oncogenic signaling and suggest that chromosome-scale dosage changes represent a fundamental route through which cancer cells optimize their evolutionary fitness.

## Methods

### Organoid lines

HUB-02-A2-130 (HUB130) healthy human colon organoids were obtained from the Hubrecht Organoid Technology (HUB) biobank under a Material Transfer Agreement. C002N healthy human colon organoids were established under previously approved protocols (B15PON) and obtained from the Netherlands Cancer Institute biobank under a Material Transfer Agreement and Institutional Review Board approval (IRBm22-310). P9T colorectal cancer organoids were obtained from a previously established biobank^41,42^. *APC* mutations were introduced into HUB130 and C002N organoids using CRISPR-Cas9 genome editing (see below for details).

### Organoid culture

Organoids were embedded in 10 µL droplets of Cultrex RGF BME Type 2 (Bio-Techne) and plated in pre-warmed culture plates. Cultures were maintained at 37°C under 5% CO2. For passaging, organoid–BME droplets were mechanically disrupted by vigorous pipetting in 1 mL ice-cold base medium consisting of Advanced DMEM/F12 (Thermo Fisher Scientific) supplemented with 10 mM HEPES (Invitrogen), 2 mM GlutaMAX (Invitrogen), and 100 units/mL penicillin–0.1 mg/mL streptomycin (Thermo Fisher Scientific). Organoid suspensions were collected to 15 mL tubes and washed with 10 mL ice-cold base medium to remove residual BME. Pelleted organoids were then dissociated into single cells by incubation in TrypLE Express (Thermo Fisher Scientific) at 37°C, resuspended in ice-cold BME and replated in pre-warmed culture plates. HUB130 and C002N wildtype organoids were cultured in base medium supplemented with 1× B27 (ThermoFisher Scientific), 1.25 mM N-acetylcysteine (Sigma-Aldrich), 500 nM A83-01 (Tocris), 50 ng/mL hEGF (PeproTech), 100 ng/mL hIGF-1 (BioLegend), 50 ng/mL hFGF-2 (PeproTech), 100 µg/mL primocin (Bio-Connect), 0.5% WNT-surrogate conditioned medium (in-house), 10% Noggin conditioned medium (in-house), and 20% R-spondin1 conditioned medium (in-house). WNT-surrogate conditioned medium was omitted and replaced with base medium for all culturing of *APC*-mutant organoids, and for all experiments with wildtype organoids where CHIR99021 (Bio-Techne) was added. P9T organoids were maintained in base medium supplemented with 1× B27 (ThermoFisher Scientific), 10 mM nicotinamide (Sigma-Aldrich), 1.25 mM N-acetylcysteine (Sigma-Aldrich), 50 ng/mL EGF (PeproTech), 500 nM A83-01, 10 µM SB202190 (Sigma-Aldrich), 100 µg/mL primocin (Bio-Connect), 10% Noggin conditioned medium (in-house), and 10% R-spondin1 conditioned medium (in-house). For all organoid lines, 10 µM Y-27632 (Merck Life Science) was added to the culture medium for the first 48 hours after plating, and medium was refreshed every 2–3 days. Cultures were routinely tested and confirmed to be negative for mycoplasma contamination.

### Organoid electroporation

Penicillin-streptomycin and primocin were removed from the culture media 4 days prior to electroporation, 10 µM Y-27632 was added to the culture media 48 hours prior to electroporation, and 1.25% (vol/vol) DMSO was added to the culture media 24 hours prior to electroporation. Organoids were dissociated in TrypLE express until small clusters of 2-5 cells remained, washed in Opti-MEM (Thermo Fisher Scientific) and resuspended in 100 µL ice-cold BTXpress electroporation buffer (Thermo Fisher Scientific). Organoid fragments were mixed with 10 µg endotoxin-free plasmid DNA (Qiagen EndoFree Plasmid Maxi kit) in 2-mm gap electroporation cuvettes (Sonidel, EC002S) and electroporated with a NEPA21 electroporator with settings as reported previously^80^. Following electroporation, cells were incubated for 30 min at room temperature before replating in BME. Organoids were switched back to regular culture medium 4 days after electroporation.

### Generation of APC mutant organoid lines

The Cas9-puro-APC_gRNA CRISPR targeting vector was a kind gift from Madelon Maurice (UMC Utrecht, the Netherlands). The gRNA encoded in this plasmid (5’-TACATCTGCTAAACATGAG-3’) was previously described^30,81^, and targets the mutation cluster region of *APC*. 10 µg plasmid DNA was electroporated into HUB130 or C002N wildtype organoids as described above. Nine days post-electroporation, organoids were passaged and selection was initiated by withdrawal of WNT-surrogate and RSPO1 from the culture medium. Clonal organoid lines were derived from the surviving population by single-cell seeding. Briefly, organoids were dissociated to single cells, filtered through a 40 µm cell strainer (pluriSelect), plated in BME, and expanded for 7 days. Individual organoids were manually picked by pipetting, seeded into separate wells, and expanded for 2 weeks prior to further passaging. After sufficient expansion had taken place, genomic DNA was isolated using the QIAmp DNA micro kit (Qiagen). The targeted *APC* locus was amplified by genomic PCR (FW: 5’-GAGAAACGTCATGTGGATCAGC-3’, REV: 5’-ACAGGTGGAGGTAATTTTGAAGC-3’). PCR products were gel-purified, cloned into a pJET vector backbone, and transformed into competent DH5α cells. For each clone, at least six bacterial colonies were selected for colony PCR (primers FW: 5-‘ CGACTCACTATAGGGAGAGCGGC-3’, REV: 5’-AAGAACATCGATTTTCCATGGCAG-3’). Colony PCR products were gel-purified and analyzed by Sanger sequencing to confirm *APC* mutations.

### Generation of chr7-iCASP8 P9T organoid lines

To generate the iCASP8 negative selection-donor, we assembled a plasmid containing TRE-myr-DmrB-Casp8-T2A-eGFP and hPGK-T2A-rtTa-WPRE cassettes between homology arms targeting the 7q34 safe harbor locus site 257^43^. Left and right homology arms (LHA, 648 bp; RHA, 675 bp) were synthesized as gBlocks and PCR-amplified to introduce recognition sites for a non-targeting sgRNA (nt-sgRNA) flanking each arm, enabling intracellular linearization of the donor upon expression of the nt-sgRNA from the same plasmid. The homology arms were cloned into a pACEBac1 plasmid derived from PX458 and described previously^82^, from which the SpCas9–T2A–GFP cassette had been removed. The nt-sgRNA sequence was inserted into the hU6-driven sgRNA cassette to direct cleavage at the sites flanking the homology arms. The myr-DmrB–Casp8–T2A–eGFP fragment was amplified from pBabe-Fv-hCaspase8-2A-GFP (Addgene plasmid #82712)^83^ and cloned between the LHA and RHA. The hPGK–PuroR–T2A–rtTA–WPRE cassette was amplified from pCW9-tBID (kindly provided by Geert Kops, Hubrecht Institute, Utrecht, the Netherlands) and inserted between eGFP and the RHA. The TRE promoter was amplified from the same plasmid and inserted upstream of myr-DmrB–Casp8–T2A–eGFP.

For targeted integration, a pACEBac1-based SpCas9–sgRNA plasmid derived from PX459^82^ was modified by introducing the D10A mutation into SpCas9, removing the T2A–PuroR sequence and replacing the original sgRNA with an sgRNA targeting safe harbor locus site 257 (5’-GAAGCTTATATAGCACCCTG-3’)^43^.

P9T organoids were electroporated with 20 µg DNA (1:1 ratio of gRNA/Cas9(D10A) plasmid and repair template plasmid) as described above (see “Organoid electroporation”). Selection with 2 µg/mL puromycin (Merck Life Science) was initiated 7 days after electroporation. Clonal organoid lines were derived from the surviving population by single-cell seeding. Briefly, organoids were dissociated to single cells, filtered through a 40 µm cell strainer (pluriSelect), plated in BME, and expanded for 7 days. Individual organoids were manually picked by pipetting, seeded into separate wells, and expanded for 2 weeks prior to further passaging. After sufficient expansion had taken place, genomic DNA was isolated using the QIAmp DNA micro kit (Qiagen). The presence of iCASP8-eGFP-puro in the targeted safe harbor locus on chromosome 7 was verified by genomic PCR, by a combination of primers binding in the genomic DNA flanking the homology arms and inside the puromycin resistance cassette (Primer pair 1: FW = 5’-TTTGCATTGCCACATCCACAAC-3’, REV = 5’-GTGGGCTTGTACTCGGTCAT-3’, primer pair 2: FW = 5’-GCAACCTCCCCTTCTACGAG-3’, REV = 5’-AATGAGGTAATTCTCCTGAGCG-3’) (Fig S5A). Additionally, based on the higher PCR-efficiency of short amplicons compared to longer amplicons, the chromosome 7 5’ forward and 3’ reverse primer were combined to check for the presence of at least one untargeted allele (Fig S5A).

To identify clones with a single allele targeted, qPCR was performed using FastStart SYBR Green Master mix (Roche), using 1 ng of genomic DNA as template. Each clone was checked by one primer pair binding to the naive version of the allele (FW: 5’-CTTCTTCTGAGGGCTGCTCC-3’, REV: 5’-ATACCCATGCAAGCCCACAA-3’) and one primer pair binding to the iCASP8-T2A-eGFP-Puro integration (iCASP8 FW: 5’-GAGCCTGAGAGAGCGATGTC-3’, T2A REV: GGATTCTCCTCCACGTCACC) (Fig S5C, D). Relative DNA levels were determined using the ΔΔCt method by normalization to an untargeted region on chromosome 7 (FW: 5’-GGAAAATCCCAGGTCCCAGG-3’, REV: 5’-TCCAAATGAGACTGCCCCAC-3’). The number of integrations was determined by normalizing each clone to the parental organoid line harboring 3 untargeted alleles, or a clone for which the presence of 2 targeted alleles could be inferred from other experiments (clone 12: absence of untargeted allele in genomic PCR, and loss of 7q-arm fragment including iCASP8-integration site based on lpWGS) (Fig S5A, B).

### Single cell karyo-sequencing (scKaryo-seq)

For scKaryo-seq, 150 µL organoid–BME droplets were mechanically disrupted by vigorous pipetting in 1 mL ice-cold base medium consisting of Advanced DMEM/F12 (Thermo Fisher Scientific) supplemented with 10 mM HEPES (Invitrogen), 2 mM GlutaMAX (Invitrogen), and 100 U/mL penicillin and 0.1 mg/mL streptomycin (Thermo Fisher Scientific). Organoid suspensions were transferred to 15 mL tubes and washed with 10 mL ice-cold base medium to remove residual BME. Organoids were subsequently dissociated to single cells by incubation with 1 mL TrypLE (Thermo Fisher Scientific) at 37°C. The resulting cell suspension was diluted to 10 mL in ice-cold base medium, filtered through a 40-µm cell strainer (pluriSelect) and washed with PBS. 500.000 cells per condition were transferred to low-binding Eppendorf tubes and resuspended in 125 µL PBSO. Cells were fixed by adding 375 µL ice-cold 100% ethanol with incubation at −20°C for 1 h. Samples were centrifuged at 600 × g for 3 min, the supernatant was removed, and the cell pellet was resuspended in 500 µL freeze buffer with the following composition: 1x cOmplete™, EDTA-free Protease Inhibitor Cocktail (Sigma-Aldrich), 0.02M HEPES pH 7.5 (Fisher Scientific), 0.1M NaCl (Invitrogen), 0.0072% Spermidine (Sigma-Aldrich), 0.1% Tween (Sigma-Aldrich), 0.002M EDTA (Invitrogen), 10% vol/vol DMSO (Invitrogen) in Ambion Nuclease Free Water (Fisher Scientific). Fixed cells were spun down at 600 rpm for 3 minutes, washed twice with PBS, and stained with 1 μg/ml DAPI diluted in PBS for 15 min. Based on the DAPI intensities, single G1 cells were sorted into 384-well plates containing 10 μl of mineral oil per well (Sigma-Aldrich). Samples were further processed as described previously^16^. Single cell libraries were sequenced on a NextSeq2000 2×100 bp sequencer, at a coverage of 16 million reads per 384 single cells. The fastq files were mapped to GRCh38 using Burrows–Wheeler aligner. For copy number analysis, aligned read data (.bam files) were analyzed in R-studio (R version 4.2.1) with the Aneufinder() function of the AneuFinder package (version 1.26.0)^84^ using the following settings: binsizes = 1e+06, stepsizes = 5e+05, remove.duplicate.reads = TRUE, min.mapq = 10, correction.method = “GC”, method = c(“dnacopy”, “edivisive”) and a euploid reference^85^. To classify cells as diploid or aneuploid, and aneuploidy as structural or numerical, we used the collection of R scripts available from the AneuFinderDataFunctions GitHub repository (https://github.com/TWvR/AneufinderDataFunctions).

### *In vitro* evolution

Organoids were treated with 50 nM GSK923295 (Bio-Connect) and 800 nM NMS-P715 (Bio-Connect) for 24 hours. Drug concentrations were determined empirically by live-cell imaging, titrating NMS-P715 to the minimum dose that prevented mitotic arrest or delayed mitosis in the presence of GSK923295. Following treatment, organoids were dissociated to single cells in TrypLE at 37°C, resuspended in BME, and plated in pre-warmed culture plates. Organoids were subsequently propagated for 12 passages as described above (see “Organoid culture”). Every third passage, an aliquot of cells was used for DNA isolation and cryopreservation in CELLBANKER I (AmsBio, #11910). To determine arm-level copy number alterations, genomic DNA was subjected to low pass whole genome sequencing and AneuFinder analysis as described below (see “Low pass-whole genome sequencing”). For each sample, arm-level copy number was calculated as the mean copy number across all bins assigned to each arm, with chromosome arm boundaries defined using centromere positions from the UCSC hg38 cytoBand annotations. Bins spanning the centromere were excluded from arm-level summaries. Chromosome arms were considered aneuploid if their mean copy number was >2.1 or <1.9.

### Low pass-whole genome sequencing (lpWGS)

Genomic DNA was isolated using the QIAamp DNA micro kit (Qiagen), with an additional 10 minute incubation with 1mg/mL RNAse A at 37°C following lysis. DNA concentration was determined using the Qubit dsDNA HS Assay Kit (Invitrogen). Low pass-whole genome sequencing was outsourced to Genewiz (Azenta Life Sciences). For copy number analysis, aligned read data (.bam files) were analyzed in R-studio (R version 4.2.1) with the Aneufinder() function of the AneuFinder package (version 1.26.0)^84^ using the following settings: binsizes = 1e+06, stepsizes = 5e+05, remove.duplicate.reads = TRUE, min.mapq = 10, correction.method = “GC”, method = c(“dnacopy”, “edivisive”) and a euploid reference^85^.

### 15x whole genome sequencing

Genomic DNA was isolated using the QIAamp DNA micro kit (Qiagen), with an additional 10 minute incubation with 1 mg/mL RNAse A at 37°C following lysis. DNA concentration and purity were determined using the Qubit dsDNA HS Assay Kit (Invitrogen). Library prep and 15x whole genome sequencing were provided by USEQ, Utrecht, the Netherlands and performed on the Illumina NovaSeq X platform using a 10B flow cell with a 300-cycle configuration (2×150 bp). FASTQ files of matched wildtype-*APC* mutant or diploid-aneuploid organoid lines were used as input for the Hartwig Medical Foundation analysis pipeline (https://github.com/HartwigMedical/pipeline5) using GRCh38 as human reference genome as described previously^86^. Variant interpretation and classification (pathogenic, likely pathogenic, benign, likely benign, or variant of uncertain significance/VUS) were performed by a certified Clinical Molecular Biologist in Pathology (KMBP; Dutch registration).

### Flow cytometry

For EdU incorporation assays, organoids were incubated in fresh media containing 10 µM EdU (Invitrogen) for 5.5 hours at 5 days post-single cell plating. Fifteen minutes prior to harvesting, 1 mg/ml Dispase II (Life Technologies) was added to the culture medium. Single cells from organoids were collected by TrypLE incubation at 37°C, filtered through a 0.4 µm pluristrainer (pluriSelect) into a 15 mL falcon tube, and washed with 10 mL PBS. Fixation, permeabilization and staining were performed using the Click-iT™ Plus EdU Alexa Fluor™ 488 Flow Cytometry Assay Kit (Invitrogen) according to the manufacturer’s protocol. Flow cytometry was performed using a FACS Celesta Cell Analyzer (BD Biosciences) with FACSDiva software (BD Biosciences).

### Reverse transcription (RT)-qPCR

Organoids were cultured in 12-well plates until near-confluency. Treatments were performed as described. Following a 15 minute incubation with 1 mg/mL Dispase II (Life Technologies), organoids were harvested into 15 mL falcon tubes and washed two times: once in ice-cold base medium and once in ice-cold PBS. RNA was isolated from organoid pellets using the RNeasy Mini Kit (Qiagen). For production of cDNA, RNA was reverse transcribed using the iScript cDNA synthesis Kit (BioRad). qPCR was performed using IQ SYBR green mix (BioRad) according to the manufacturer’s protocol. Gene expression was calculated using the ΔΔCt method with normalization of expression values to reference gene *HPRT1*. Table 1 indicates RT-qPCR primers.

**Table 1.**
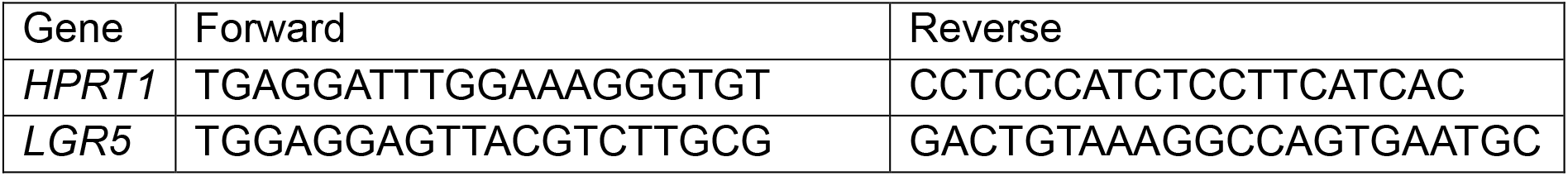
RTqPCR primer sequences.

### IncuCyte-based growth assessment

Organoids were seeded as single cells in 96-well plates (Corning) at a density of 3500 cells in 5 µL BME per well. Organoid growth was monitored by imaging every 8 or 12 hours using an IncuCyte system with an organoid software analysis module (Sartorius). Total organoid area was used as measure for growth. All growth curves were normalized to the same early timepoint (t=8h or t=12 hour, depending on experiment). For plots where only one time point is displayed (Figure panels: 5B, C, E, F, G, 6B), the final timepoint with accurate organoid segmentation was selected per experiment (∼120 hours).

### Quantification of viable cells by CellTiterGlo

Organoids were seeded as single cells in white clear-bottom 96-well plates at a density of 3500 cells in 5 µL BME per well. Media was refreshed every 2-3 days. At day 6 after seeding, organoid viability was quantified using the CellTiter-Glo 3D Cell Viability Assay kit (Promega) according to manufacturer’s instructions. Luminescence was quantified using a Centro LB 960 Microplate Luminometer (Berthold Technologies)

### CASP8 induction and recovery of CASP8-activation survivors

Organoids were dissociated to single cells by incubation with TrypLE and filtered through a 40-µm PluriStrainer (pluriSelect) to ensure a single-cell suspension. Single cells were plated in BME in pre-warmed 24-well plates in regular growth medium containing 10 µM Y-27632, 1 µg/ml doxycycline to induce CASP8 expression and 10 nM AP20187 (Merck) to induce CASP8 dimerization. Control organoids were seeded in parallel in medium containing 10 µM Y-27632 and a volume equivalent of DMSO. Medium was refreshed every 2–3 days with fresh doxycycline and AP20187, or DMSO in control wells, for 7–10 days. Surviving organoids were manually picked and seeded individually in BME in pre-warmed 48-well plates. After isolation, organoids were cultured in regular medium supplemented with 10 µM Y-27632 and without doxycycline, AP20187 or DMSO. Brightfield images were acquired at the indicated time points using an EVOS M5000 Imaging System (Thermo Fisher Scientific). Organoid size was quantified from brightfield images using a custom Fiji macro. In some experiments, organoids were exposed to 50 nM CENP-E inhibitor GSK-923295 (Bio-Connect) and 500 nM MPS1 inhibitor NMS-P715 (Bio-Connect), or to an equivalent volume of DMSO, for 24 h before single-cell dissociation. This pretreatment was used as an attempt to increase the recovery of organoids that had lost a full copy of chromosome 7. However, in the P9T organoid model, this treatment did not increase the frequency of chromosome 7 loss among the sequenced organoids (note that this organoid line has high baseline missegregation frequencies^42^). Treated and untreated organoids were therefore pooled for downstream analyses.

### Noggin withdrawal or iCRT3 treatment

Organoids were treated as described above, with the following modification: concomitantly with the initiation of doxycycline and AP2081 treatment, either Noggin withdrawal or iCRT3 treatment was initiated. For Noggin withdrawal, Noggin-conditioned medium was omitted and replaced with base medium. For iCRT3 treatment, organoids were cultured in the presence of 15 µM iCRT3 (Bio-Connect). Both Noggin withdrawal and iCRT3 treatment were maintained until sample collection.

### Single organoid DNA sequencing

At 12-14 days following single organoid plating, single-cell derived poorly growing organoids were harvested in ice-cold CELLBANKER I (AmsBio, #11910) by pipetting vigorously up and down, and transferred to a 6 cm cell culture dish. Organoids were then collected by manual picking in 3 µL volume into 96-well plates and frozen at −70°C until further processing. Organoids were prepared for sequencing using the Agilent BRAVO liquid handler system. After organoid lysis, DNA was fragmented by micrococcal nuclease. Following clean up, end repair and A-tailing of the DNA fragments was performed in a single reaction mix containing T4 DNA polymerase, T4 polynucleotide kinase and Bst 2.0 warm start polymerase by first incubating 30min at 25C (end repair) and subsequently incubating 30min at 68°C (A-tailing). IDT stubby adapters were ligated to either side of the DNA fragments by addition of ligation reaction mix containing T4 DNA Quick ligase to the same tube. After clean up, the adapter-ligated DNA fragments were immediately subjected to 17 cycles of PCR using Phusion High Fidelity DNA polymerase and IDT unique dual 10bp indexed primers. A final clean-up was performed using AMPure beads, and DNA was eluted in 6 µl elution buffer. Sequencing was performed on the NextSeq 2000 instrument (Illumina) with 77 cycles single end read length, or on a NovaSeq X Plus instrument (Illumina) with a paired-end read length of 150 cycles. The generated data were demultiplexed using sample-specific barcodes and converted into fastq files using bcl2fastq. The aligned read data (bam files) was analyzed using AneuFinder software as described previously^84^.

### Bulk RNA sequencing

Three biological replicates of HUB130 APCmut C9 parental, diploid clone 2, trisomy 7 clone 3 and trisomy 7 clone 4 were harvested and snap-frozen in liquid nitrogen at day 5 after single cell seeding. RNA sequencing was provided by USEQ, Utrecht, the Netherlands. RNA isolation was performed on a QIAsymphony (Qiagen) using the QIAsymphony RNA Kit (Qiagen) and miRNA_CT_400 protocol (Qiagen). RNA libraries were prepared using a TruSeq Stranded mRNA Preparation Kit (Illumina). Sequencing was performed on the Illumina NovaSeq X platform using a 10B flow cell with a 300-cycle configuration, yielding 150 base pair paired-end reads (2×150 bp). Quality control on the sequence reads from the raw FASTQ files was done with FastQC (v0.11.9). TrimGalore (v0.6.7) was used to trim reads based on quality and adapter presence after which FastQC was again used to check the resulting quality. rRNA reads were filtered out using SortMeRNA (v4.3.6) after which the resulting reads were aligned to the GRCh38 reference genome using the STAR (v2.7.10b) aligner. Read counts were then generated using the Subread FeatureCounts module (v2.0.3) with the Homo_sapiens.GRCh38.106.ncbi.gtf file as annotation. Differential expression analysis was performed using DESeq2^87^ (v1.38.0).

### Single organoid RNA sequencing

#### Sample collection

P9T chr7-iCASP8 C9 organoids were treated as described under **‘**CASP8 induction and recovery of CASP8-activation survivors’. At 12–14 days after single-organoid seeding, growth-impaired dox/AP-surviving organoids and size-matched control organoids were harvested in ice-cold PBS by vigorous pipetting to dissociate the BME and transferred to a 6-cm cell-culture dish. Individual organoids were manually collected in 2.5 µL and immediately snap-frozen in liquid nitrogen.

#### RNA sequencing

P9T RNA-sequencing libraries were prepared using the Smart-seq3 protocol^88^. Paired-end sequencing was performed with forward reads containing a 0-, 3-, or 6-nucleotide variable spacer followed by a GGG motif and the cDNA-derived transcript sequence, while reverse reads contained the unique molecular identifier (UMI). Sequencing was performed on a NextSeq 2000 instrument (Illumina) with a single-end read length of 77 cycles. Paired-end reads were processed using the nf-core/rnaseq pipeline v3.15.1. Six-nucleotide UMIs were extracted from read 2 using UMI-tools, followed by removal of read 2 and UMI-aware deduplication. Reads were aligned and quantified with STAR and Salmon against the GRCh38 primary assembly using Ensembl release 109 gene annotations. Quality-control metrics, including UMI deduplication statistics, were summarized with MultiQC.

#### Validation of RNA-based copy number inference

RNA-derived copy-number profiles were validated using matched bulk RNA-sequencing and low-pass whole-genome sequencing (lpWGS) from the P9T chr7-iCASP8 C9 line, which carries three copies of chromosome 7, and a derivative of this line that had previously lost the third copy of chromosome 7 following dox/AP treatment that proliferated sufficiently for bulk DNA and RNA isolation (“dis7-resc1” in Fig S7). Gene-level expression changes were estimated relative to C9, with normalization accounting for the lpWGS-defined copy-number profile of the trisomic reference; fold changes were then centered on copy-neutral regions and smoothed across genomic position to infer chromosome-scale dosage. The inferred RNA profiles recovered the chromosome 7 alterations identified independently by lpWGS (Fig S7C). To assess the minimal required sequencing depth for correct copy number inference, we downsampled the non-reference bulk RNA-sequencing libraries with scuttle and compared mean RNA-derived fold changes within lpWGS-defined segments with the corresponding DNA copy-number changes using segment-length-weighted root mean squared error. Concordance deteriorated below approximately 25,000 gene-assigned reads, which was therefore used as the minimum threshold for RNA-based copy-number inference.

Individual organoids containing more than 25,000 gene-assigned reads were then scored against the same bulk P9T chr7-iCASP8 C9 reference. Gene-level expression fold changes were centered on the modal genome-wide signal and smoothed across genomic position within each chromosome. The resulting profiles were converted to approximate copy numbers by anchoring the chromosome-level signal of C9 organoids to the lpWGS-defined C9 copy-number profile. As controls we included derivatives of the P9T chr7-iCASP8 C9 organoid line that had previously survived dox/AP treatment and been established as clonal organoid lines with known karyotype: one line that underwent the full dox/AP procedure but retained a trisomy 7 karyotype (“tris7-ctrl”) and two lines that lost the extra copy of chromosome 7 following dox/AP treatment, without suffering from the growth arrest typically associated with trisomy 7 loss (“dis7 spontaneous rescue”). Copy number accurately reflected known karyotypes for these organoid lines (Fig S7D). Note that “dis 7 spontaneous rescue” organoids were only included as controls for RNA-based copy number inference, and were not included for differential expression analysis.

#### Differential expression analysis

RNA sequencing and RNA-based copy-number inference were performed as described above. Organoids with full trisomy 7 loss or 7q loss were manually selected and included in differential expression analysis against control organoids using the R packages *DESeq2*^87^ (version 1.46.0) and GSVA (version 2.0.7).

### Gene Ontology enrichment analysis

Differentially expressed genes were selected from the DESeq2 analysis using baseMean ≥ 1, adjusted P < 0.05 and an absolute log2 fold-change threshold of 0.58. Upregulated and downregulated genes were analyzed separately for enrichment of Gene Ontology Biological Process terms using g:Profiler with an unordered query. The statistical domain was restricted to annotated genes, and terms containing between 10 and 500 genes were retained. Significance was assessed using the Benjamini–Hochberg false-discovery-rate correction with a threshold of 0.05.

### Gene expression analysis in patient CRCs

TCGA-COADREAD primary-tumor RNA-sequencing, somatic mutation, clinical, MSI, and allele-specific copy-number data were obtained from the NCI Genomic Data Commons^2,89^. For each chromosome, segments were classified as gain, neutral, or loss relative to the tumor’s modal integer copy number calculated from the other autosomes. An arm was considered gained or neutral when at least 80% of its length had the corresponding state. Whole-chromosome 7 gain was defined by gain of both 7p and 7q, whereas the reference group required both arms to be neutral. Tumors with losses, discordant arms, or mixed states were excluded.

APC alteration was inferred from nonsynonymous APC mutations together with deletion or loss of heterozygosity at the APC locus and was coded as altered versus unaltered. Whole-genome duplication was defined by a major-allele copy number of at least two across at least half of the measured autosomal genome. TP53 mutation and RAS–RAF mutation indicated nonsynonymous mutations in TP53 or in KRAS, NRAS, or BRAF, respectively. Tumor site was classified as left colon, right colon, rectum, or other. The analysis was restricted to MSS tumors and excluded tumors carrying nonsynonymous POLE mutations.

Raw RNA counts were analyzed using R package DESeq2^87^ (version 1.46.0). The fully adjusted model included whole-chromosome 7 gain, APC alteration, whole-genome duplication, TP53 mutation, RAS–RAF mutation, and tumor site. Wald standard errors were used to calculate 95% confidence intervals. Within each prespecified gene panel, gene-level Wald (P)-values were combined using Fisher’s method.

### Chromosome 7 gain and APC inactivation level analysis in patient CRCs

Primary, microsatellite-stable (MSS) colorectal adenocarcinomas with biallelic *APC* mutations were analyzed from TCGA-COADREAD (n = 214) and the 100,000 Genomes Project (100kGP, n = 829) CRC cohorts.

Biallelic *APC* status and total retained 20-amino-acid repeats (20Rs) were determined as in Brunet Guasch et al.^55^ using genomic annotations reported by Cornish et al.^1^ Given copy number at the *APC* site [a, b] (“a” for major allele and “b” for the minor allele), samples with no WGD were classified at initiation as diploid if [a>0, b>0], copy-loss loss of heterozygosity (CL-LOH) if [1,0], and copy-neutral loss of heterozygosity (CN-LOH) if [a ≥ 2,0]. Under the rationale that WGD occurred after *APC* inactivation, samples with WGD were classified as diploid at initiation if [a > 0, b > 0], CL-LOH if [1–2, 0], or CN-LOH if [a > 2,0]. Tumors with two truncating mutations were assumed to have inactivation in each allele, and to retain the sum of the number of 20R domains upstream of each mutation. Tumors with a single mutation and no informative copy-number evidence were excluded. In the TCGA cohort, allele-specific copy number was not available, and discrete copy-number calls at the *APC* locus were instead obtained from cBioPortal. Diploid tumors with two truncating mutations were assigned a total retained 20Rs as the sum of the 20R domains upstream of each mutation. For tumors with a single truncating mutation, only a heterozygous loss (GISTIC −1) call was treated as informative, and was interpreted as equivalent to CL-LOH, contributing 0 retained repeats.

Tumors were classified as chromosome 7 gain if both 7p and 7q showed copy-number gain, or without gain if both arms were unaltered. Tumors were also classified as mutant for *AMER1*, *BCL9*, *SOX9*, or *TCF7L2* if carrying an annotated oncogenic mutation in the respective gene, or wild-type otherwise; these genes have been reported to act as secondary modifiers of WNT activity in combination with APC loss^55^. This comparison was restricted to primary MSS tumors in the 100kGP cohort, and the difference in mean total retained 20Rs between mutant and wild-type tumors was calculated for each gene as in the chromosome 7 comparison above.

## Author contributions

Conceptualization - ICZ, PCG and SMAL; methodology - ICZ, PCG, MBG, MDN, FF, MS and SMAL; investigation – ICZ, PCG, MV, NM, AV, MBG, MS and SMAL; formal analysis - ICZ, PCG, RW, DK SM, MS, SMAL; visualization - ICZ, PCG, MBG, SM, HS, MS; resources – BC, HS; funding acquisition – SMAL; project administration – SMAL; writing original draft – ICZ, PCG, SMAL; writing review & editing – ICZ, SMAL; supervision MDN, FF, SMAL.

## Acknowledgements

We thank Anne Jansen for variant interpretation; Ingrid Jordens for support with single cell sorting and useful suggestions; Michael Hadders, Jelte van der Vaart and the Aniek Janssen lab for useful suggestions; Madelon Maurice and Thomas van Ravesteyn for critical reading of the manuscript; Theofilos Chalkiadakis and Flip Mulder for bioinformatics support. We acknowledge Single-Cell Core, Utrecht, the Netherlands for performing the single-cell sample preparation and we acknowledge the Utrecht Sequencing Facility (USEQ) for providing sequencing service and data. We gratefully acknowledge the participants of the National Genomic Research Library (NGRL), whose contributions made this research possible. Secure access to the NGRL under project ID 26 was provided by Genomics England, which delivers the NGRL in partnership with NHS England, and is wholly owned by the UK Department of Health and Social Care. The NGRL contains participants’ health data collected by the NHS as part of their care, along with samples and data from their participation in research, for which fully informed consent has been obtained. This includes genomic and clinical data provided through the NHS Genomic Medicine Service, as well as data obtained through research studies, including the 100,000 Genomes Project and the Generation Study, both of which are delivered in partnership with the NHS, and from other research cohorts involving external collaborators.

This work was supported by the Dutch Cancer Society (KWF) research grant 13515, awarded to SMAL. This work is also part of Oncode Institute (ICZ, PCG, MV, NM, AV, SM, HJS, and SMAL), which is partly financed by the Dutch Cancer Society. Work in the FF lab is supported by a ZonMw Vici grant (09150182210049). iPsomics is funded by a ZonMw/PharmaNL grant. Work in the HJS lab is supported by an ERC Consolidator Grant (101125393). MBG is supported by the Centre for Doctoral Training in Mathematical Modelling, Analysis and Computation, University of Edinburgh (GB) (Grant No. EP/S023291/1). SM is supported by a NWO Veni grant (VI.Veni.222.187). USEQ is subsidized by the University Medical Center Utrecht and The Netherlands X-omics Initiative (NWO project 184.034.019).

## Declarations

During the preparation of this work, the authors used generative AI in order to improve readability and language of the manuscript. The author(s) reviewed and edited the content as needed and take full responsibility for the content of the published article.

BC has a patent issued on Progression markers for colorectal cancer (EP19187894) and has several patents pending or issued that are not related to this work. FF is chief scientific officer of iPsomics B.V.. He does not gain financial benefit from this role.

## Data & code availability

Code related to copy number-inference from single organoid RNA-sequencing can be found at https://github.com/mschubert/CRC-trisomy7-organoids-public

## Supplemental material

## Supplemental figures

**Figure S1:**
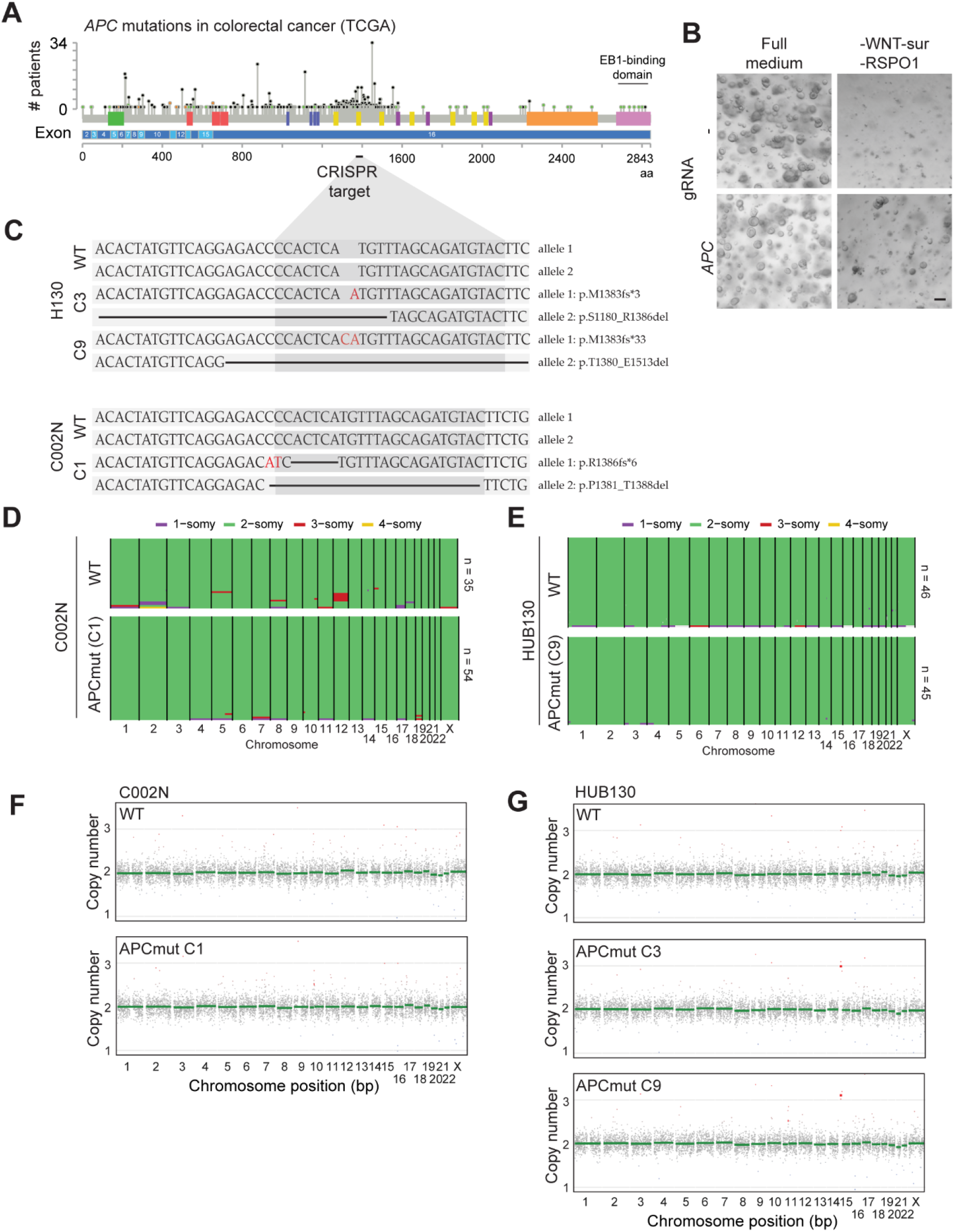
Generation and characterization of APC mutant organoids. A) Distribution of APC mutations in human colorectal cancer (TCGA). B) Representative images of mock-electroporated and APC-targeted organoids cultured in complete medium or medium lacking WNT surrogate and R-spondin1. Scale bar, 200 µm. C) Sanger sequencing of APC-wildtype and APC-mutant organoid lines. D) Single cell karyotype-sequencing of C002N wildtype and APC-mutant organoid lines. E) Single cell karyotype-sequencing of HUB130 wildtype and APC-mutant organoid lines. F) Bulk lpWGS profiles of C002N wildtype and APC-mutant organoid lines after extended passaging (≥ 12 passages). G) Bulk lpWGS profiles of HUB130 wildtype and APC-mutant organoid lines after extended passaging (≥ 12 passages).

**Figure S2:**
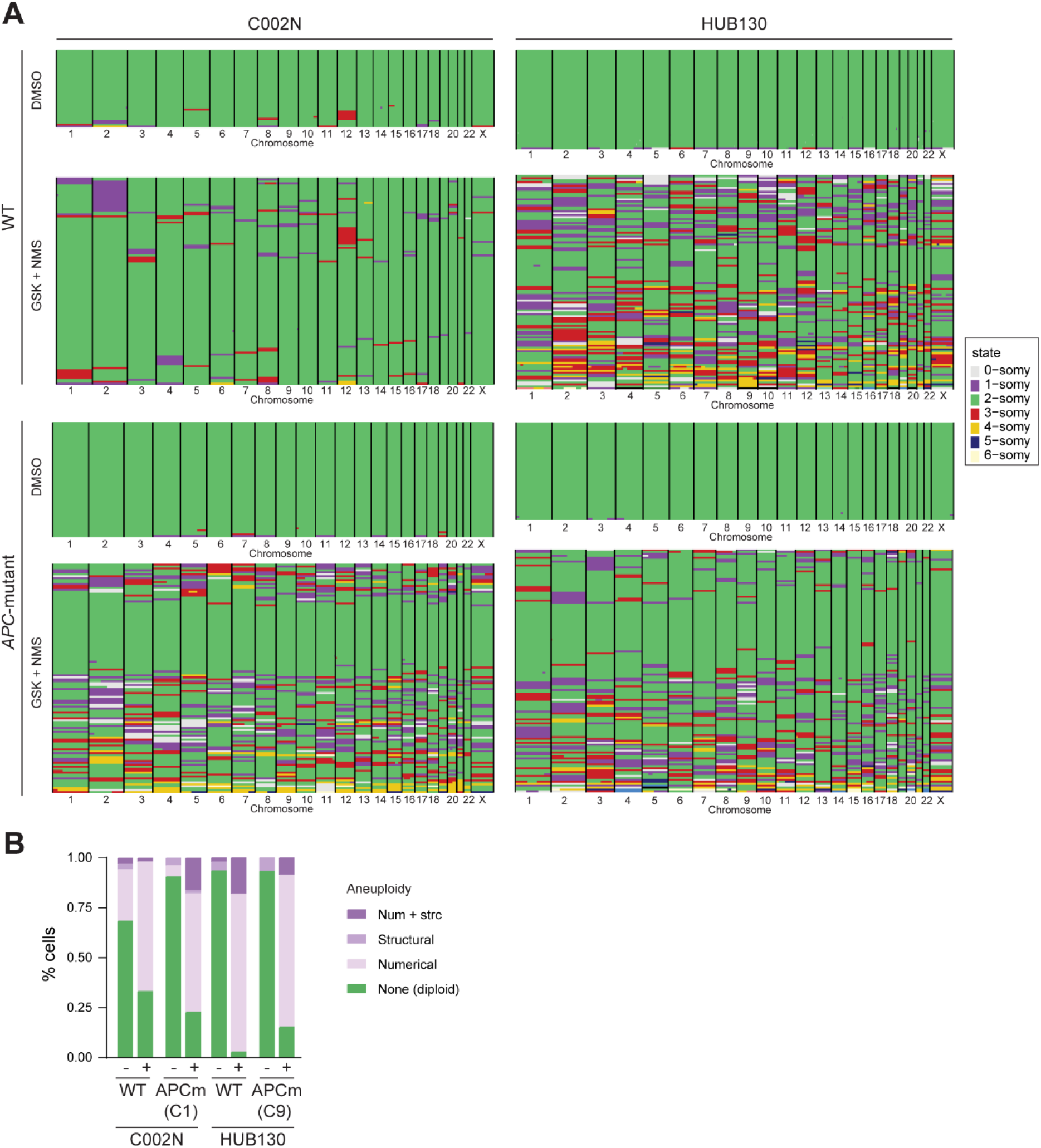
Induction of chromosomal instability using CENP-E and MPS1-inhibitors. A) Single cell karyotype-sequencing of organoids that were exposed to 50 nM GSK923295 (CENP-Ei) and 800 nM NMSp715 (MPS1i). Cell numbers per condition: C002N WT DMSO = 35, C002N WT GSK + NMS = 108, C002N APCmut DMSO = 54, C002N APCmut GSK + NMS = 118, HUB130 WT DMSO = 46, HUB130 WT GSK + NMS = 110, HUB130 APCmut DMSO = 45, HUB130 APCmut GSK + NMS = 125. B) Quantification of cells harboring chromosomes with numerical or structural aneuploidy.

**Figure S3:**
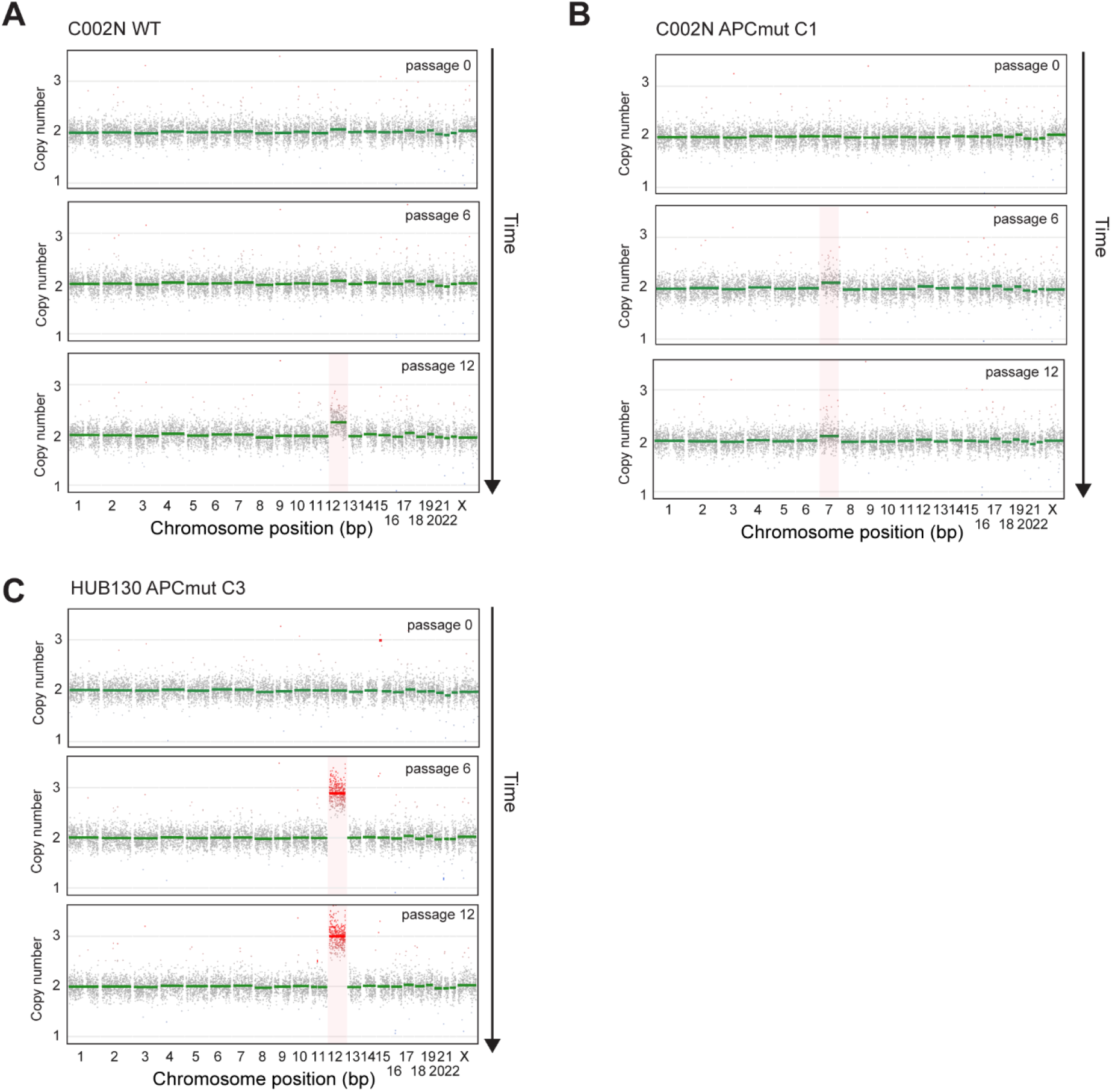
Additional examples of in vitro karyotype evolution. A-C) Representative bulk lpWGS profiles from independent in vitro evolution experiments.

**Figure S4:**
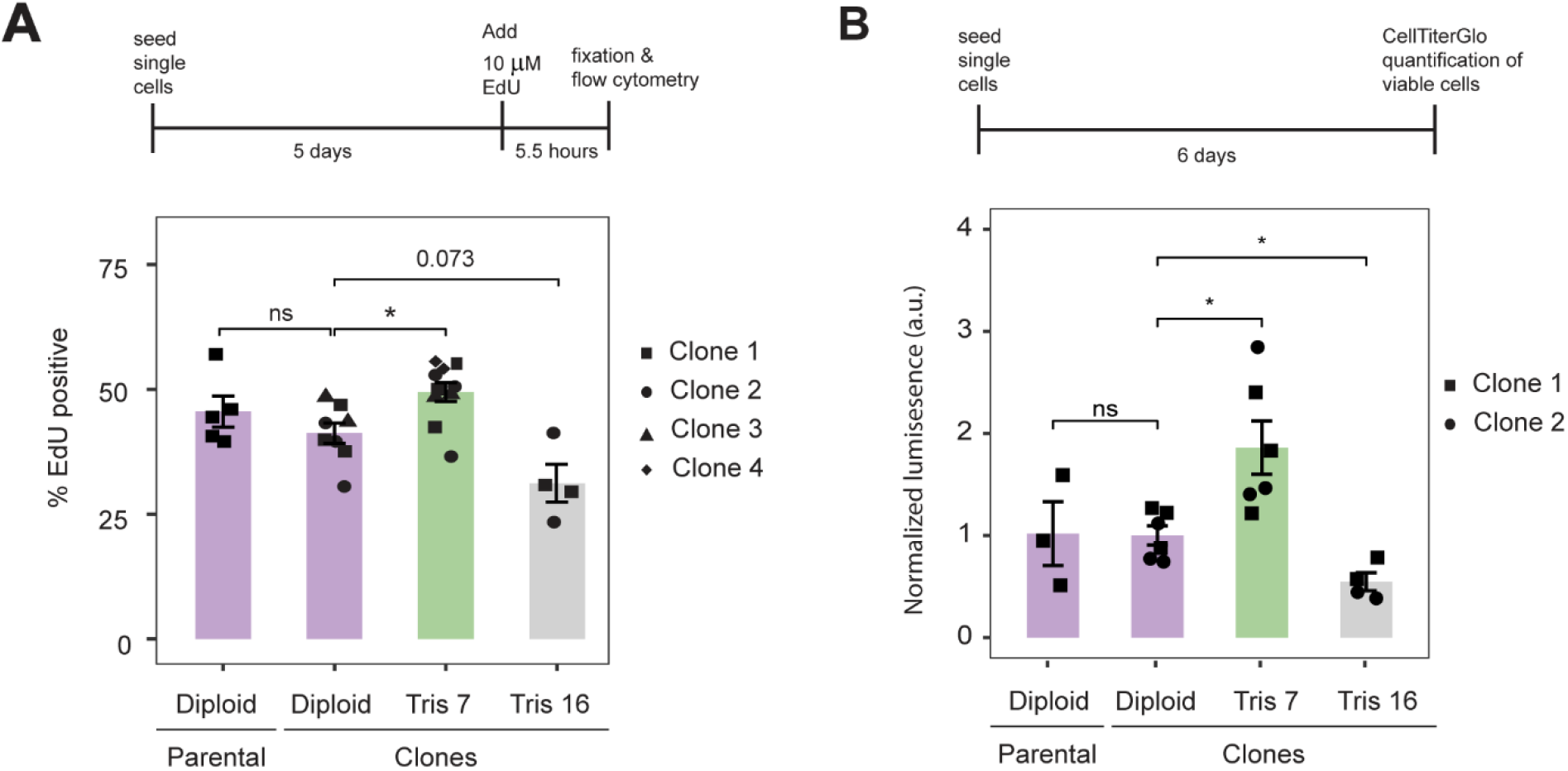
Trisomy 7 confers a cell-autonomous growth advantage. A) Percentage of EdU-positive cells after a 5.5-hour pulse labelling, quantified by flow cytometry. Mean ± SD. Dots represent biological replicates. Symbol shapes indicate independent clones. Two-sided Wilcoxon rank-sum test, * = p < 0.05. B) Normalized CellTiter-Glo luminescence 7 days after plating equal numbers of single cells. Mean ± SD. Dots represent biological replicates. Symbol shapes indicate independent clones. Two-sided Wilcoxon rank-sum test, * = p < 0.05.

**Figure S5:**
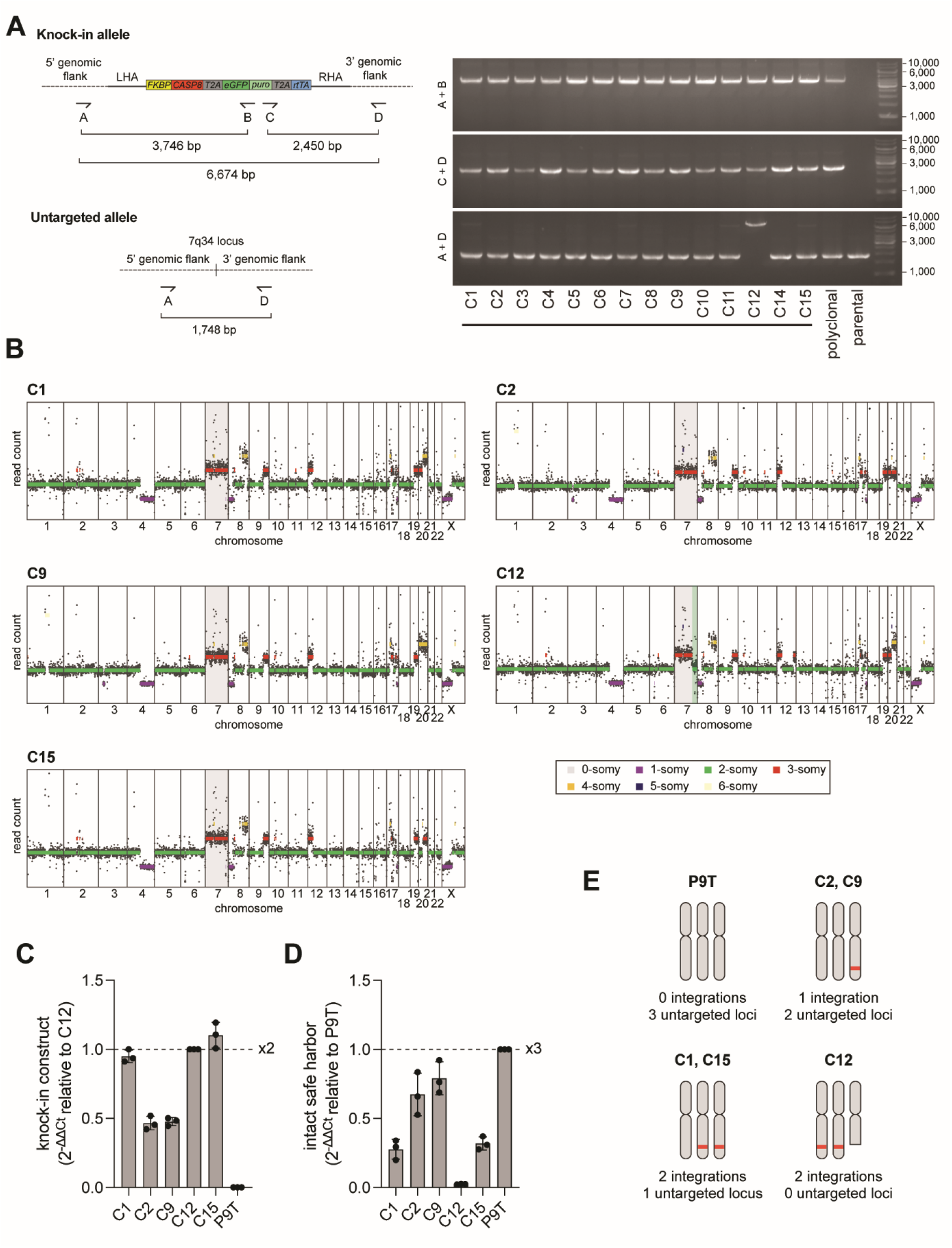
Identification of P9T chromosome 7-iCASP8 organoid clones with monoallelic cassette integration. A) Genomic PCR strategy to identify organoid clones with successful iCASP8 integration. B) Bulk lpWGS profiles P9T chr7-iCASP8 clones 1, 2, 9, 12 and 15. C) Quantitative genomic PCR measuring iCASP8 cassette copy number. Mean ± SD. Dots represent biological replicates. C12 was used as the two-copy reference, as this clone has only 2 alleles in total (figure panel B), and has no untargeted alleles (figure panel A). D) Quantitative genomic PCR of the intact safe harbor locus. Mean ± SD. Dots represent biological replicates. The parental P9T organoid line served as the reference for three intact alleles. E) Schematic summary of chr7-iCASP8 integration status across parental and clonal organoid lines.

**Figure S6:**
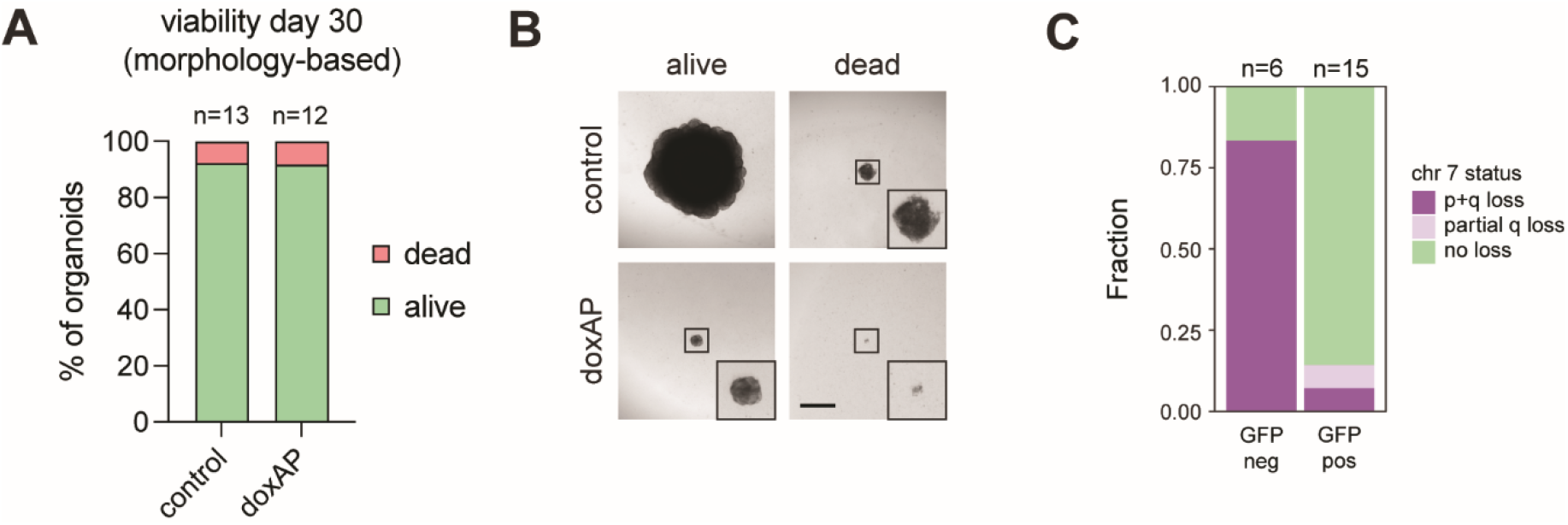
Additional characterization of organoids surviving trisomy 7 reversal. A, B) Morphology-based viability assessment of dox/AP-treated organoids. Scale bar, 500 µm. C) lpWGS of dox/AP-treated, GFP-positive organoids and GFP-negative organoids.

**Figure S7:**
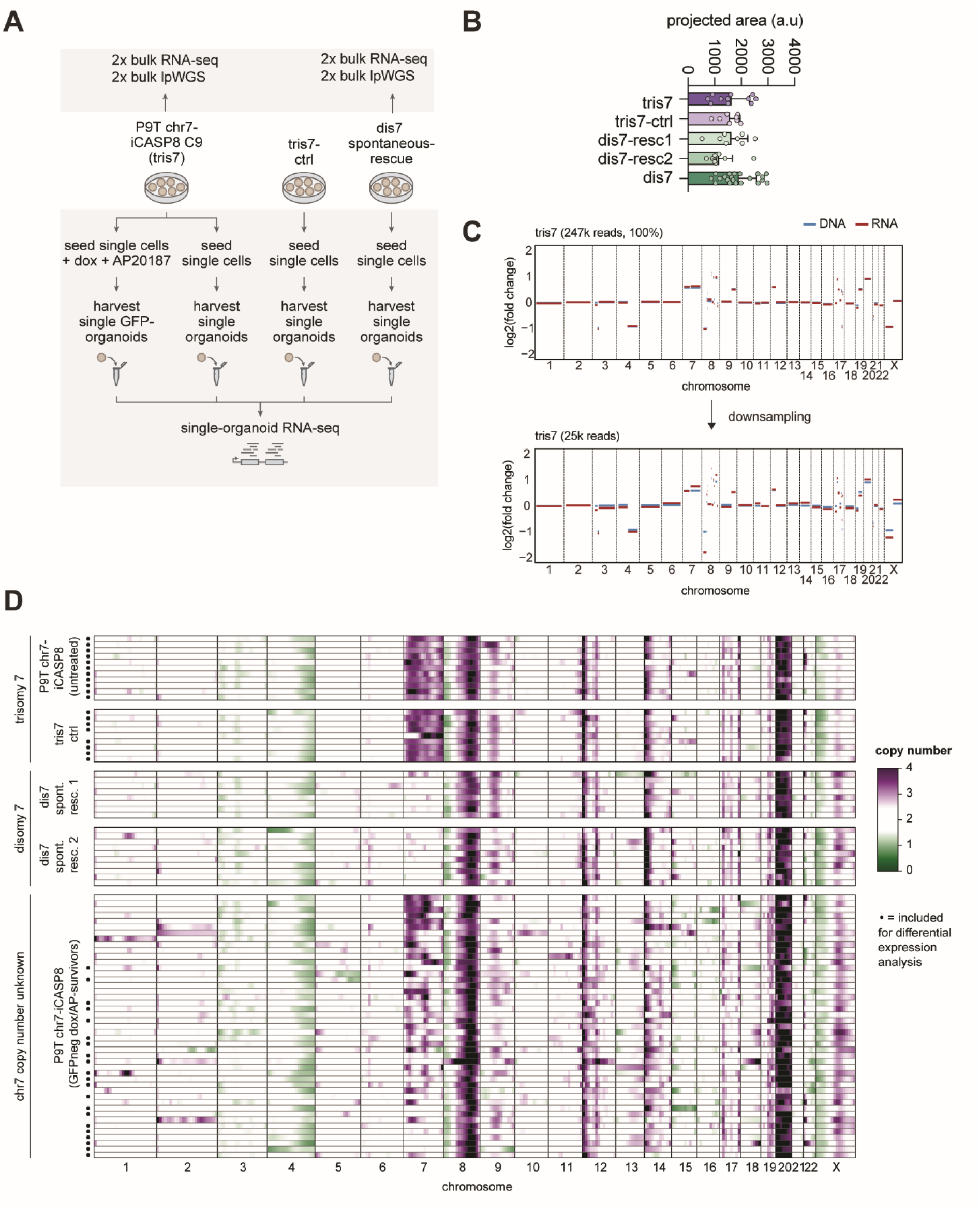
Inference of chromosomal copy number alterations from single organoid RNA sequencing. A) Schematic of experimental set-up. B) Organoid size at the time of RNA isolation. Mean ± SD. C) Validation of RNA-based copy-number inference using matched bulk lpWGS and bulk RNA-sequencing data. Reliable copy-number calling was maintained after down-sampling to 25.000 reads, which was used as the inclusion threshold for single organoids. D) Inferred chromosome copy number profiles of individual organoids. Each line represents one organoid. Dox/AP-treated organoids with loss of a full chromosome 7 or the 7q arm were manually selected for subsequent analyses (selected samples indicated with black circle).

---

**Supplemental file 1**: Overview of mutations identified in *APC* wildtype, *APC* mutant diploid, *APC* mutant trisomy 7 and *APC* mutant trisomy 16 organoid lines in 15x WGS data using the Hartwig Medical Foundation analysis pipeline.

